# Crystal structure and interaction studies of human DHTKD1 provide insight into a mitochondrial megacomplex in lysine catabolism

**DOI:** 10.1101/2020.01.20.912931

**Authors:** Gustavo A. Bezerra, William R. Foster, Henry J. Bailey, Kevin G. Hicks, Sven W. Sauer, Bianca Dimitrov, Jürgen G. Okun, Jared Rutter, Stefan Kölker, Wyatt W. Yue

**Affiliations:** Structural Genomics Consortium, Nuffield Department of Medicine, University of Oxford, UK OX3 7DQ; Department of Biochemistry, University of Utah School of Medicine; University Children’s Hospital, Division of Child Neurology and Metabolic Diseases, Heidelberg, Germany

**Author notes:** These authors contributed equally to this work (Gustavo A. Bezerra, William R. Foster). To whom correspondence could be addressed: Wyatt W. Yue.

**Keywords:** 2-oxoadipate, 2-oxoacid dehydrogenase, thiamine diphosphate, lysine catabolism, missense variants, x-ray crystallography, electron microscopy

## Abstract

DHTKD1 is a lesser-studied E1 enzyme belonging to the family of 2-oxoacid dehydrogenases. DHTKD1, in complex with the E2 (dihydrolipoamide succinyltransferase, DLST) and E3 (lipoamide dehydrogenase, DLD) components, is implicated in lysine and tryptophan catabolism by catalysing the oxidative decarboxylation of 2-oxoadipate (2OA) in the mitochondria. Here, we solved the crystal structure of human DHTKD1 at 1.9 Å resolution in binary complex with the thiamine diphosphate (ThDP) cofactor. Our structure explains the evolutionary divergence of DHTKD1 from the well-characterized homologue 2-oxoglutarate (2OG) dehydrogenase, in its preference for the larger 2OA substrate than 2OG. Inherited *DHTKD1* missense mutations cause the lysine metabolic condition 2-aminoadipic and 2-oxoadipic aciduria. Reconstruction of the missense variant proteins reveal their underlying molecular defects, which include protein destabilisation, disruption of protein-protein interactions, and alterations in the protein surface. We further generated a 5.0 Å reconstruction of the human DLST inner core by single-particle electron microscopy, revealing a 24-mer cubic architecture that serves as a scaffold for assembly of DHTKD1 and DLD. This structural study provides a starting point to develop small molecule DHTKD1 inhibitors for probing mitochondrial energy metabolism.

## INTRODUCTION

The family of multi-component 2-oxoacid dehydrogenase complexes, of which pyruvate dehydrogenase (PDHc), branched chain α□ketoacid dehydrogenase (BCKDHc), and 2-oxoglutarate dehydrogenase (OGDHc) complexes are canonical members, catalyse the oxidative decarboxylation of 2-oxoacids (e.g. pyruvate, branched-chain 2-oxoacids, and 2-oxoglutarate) into their corresponding acyl-CoA thioester, generating the reducing equivalent NADH. These biochemical reactions play crucial roles in intermediary metabolism, and are tightly regulated by phosphorylation and allosteric effectors [1,2].

The overall reaction catalysed by 2-oxoacid dehydrogenases is dissected into three sequential steps each catalysed by an individual enzyme [3,4]. In the first step, rate-limiting for the overall reaction, the E1 enzyme (a 2-oxoacid decarboxylase; EC 1.2.4.2) catalyses the irreversible decarboxylation of 2-oxoacids via the thiamine diphosphate (ThDP) cofactor and subsequent transfer of the decarboxylated acyl intermediate (reductive acylation) on an oxidised dihydrolipoyl group that is covalently amidated to the E2 enzyme (a dihydrolipoyl acyltransferase; EC 2.3.1.61). In the second step, E2 transfers the acyl moiety from the dihydrolipoyl group onto a CoA-SH acceptor, generating acyl-CoA and a reduced dihydrolipoyl group. In the final step, one FAD-dependent E3 enzyme universal to all complexes (dihydrolipoamide dehydrogenase, DLD; EC 1.8.1.4) re-oxidises the dihydrolipoyl group by transferring one reducing equivalent of NAD^+^ to yield NADH.

To achieve the overall oxidative decarboxylation reaction, multiple copies of the E1, E2 and E3 components classically assemble into a supramolecular complex reaching 4-10 MDa in weight [5]. Structural studies have shown E2 enzymes from various organisms to exist in a high-order cubic 24-mer or dodecahedral 60-mer [6], acting as a scaffold onto which copies of E1 and E3 are assembled. Such quaternary arrangement, a classic example of a metabolon, provides a means by which products of one reaction are funnelled into the catalytic centres of the next reactions to enhance enzymatic efficiency and avoid undesirable side-reactions [7]. For example, the E2-attached dihydrolipoyl cofactor is expected to shuttle catalytic intermediate substrates between the active sites of E1 and E3 enzymes by means of a ‘swinging-arm’ mechanism [8-10].

The human genome encodes five E1-type decarboxylases (PDH, BCKDH, OGDH, DHTKD1, OGDHL), among which OGDH, OGDHL and DHTKD1 form a more evolutionarily related subgroup with respect to the E1 architecture and the E2 enzyme employed [11]. The OGDHc complex, composed of OGDH as E1, dihydrolipoamide succinyltransferase (DLST) as E2, and DLD as E3, converts the metabolite 2-oxoglutarate (2OG) to succinyl-CoA and serves as a rate-limiting step in the Krebs cycle [12]. A close homologue of OGDH, DHTKD1 (dehydrogenase E1 and transketolase domain-containing protein 1) is positioned in the last step of lysine and tryptophan catabolism with the common product being 2-oxoadipate (2OA), one methylene group longer than 2OG. To catalyse the oxidative decarboxylation of 2OA to glutaryl-CoA [13], DHTKD1 recruits the same E2 (DLST) and E3 (DLD) as OGDH to form the 2-oxoadipate dehydrogenase complex (OADHc) [14,15], implying that DLST also acts as a dihydrolipoamide glutaryltransferase. DHTKD1 [16,17] joins OGDH [18,19] as a contributor of reactive oxygen species in the mitochondria via radical formation from a catalytic intermediate. DHTKD1 is increasingly recognized as essential for mitochondrial function and energy production [18,19].

In support of this role, inherited *DHTKD1* mutations are identified as the molecular cause of two rare Mendelian disorders. 2-Aminoadipic and 2-oxoadipic aciduria (OMIM 204750) is an inborn error of metabolism with questionable clinical consequence [20], characterized biochemically by increased urinary excretion of 2-oxoadipate and its transamination product 2-aminoadipate [21,22]. Among < 30 reported cases due to autosomal recessive missense and nonsense mutations [23], p.Gly729Arg and p.Arg455Gln are common variants. Additionally, a nonsense *DHTKD1* mutation causes Charcot-Marie-Tooth disease type 2Q (CMT2Q, OMIM 615025), an autosomal dominant neurodegenerative disorder characterized by motor and sensory neuropathies [24].

While structural studies have been carried out for PDH and BCKDH E1 enzymes from various organisms across the phyla, including human [25,26], only prokaryotic OGDHs have been crystallised. These include the *apo* structure of *E. coli* OGDH (ecOGDH) [27], as well as various structures of *Mycobacterium smegmatis* OGDH (msOGDH) complexed with active site catalytic intermediates [28-30]. In this study, we report the crystal structure of human DHTKD1 and a model of the human DLST catalytic core determined by cryo-electron microscopy. We also characterised disease-causing variants of DHTKD1 that destabilise the protein, or disrupt the interaction with DLST.

## RESULTS & DISCUSSION

### Overall structure of human DHTKD1 homodimer

Human DHTKD1 (hDHTKD1) is a 919-amino acid (aa) polypeptide (Fig. 1a), with the N-terminal 22 aa predicted to form the mitochondrial targeting signal peptide [11]. We expressed in *E. coli* the soluble proteins for the precursor hDHTKD1_1-919_, the predicted mature protein (hDHTKD1_23-919_), as well as a further truncated construct (hDHTKD1_45-919_) removing the putatively disordered aa 24-44 (Suppl. Fig. 1). Attempts to crystallise full-length protein were not successful despite intensive screening of conditions; however, crystals were obtained for hDHTKD1_45-919_ in the presence of both ThDP and Mg^2+^. This protein construct is active *in vitro*, exhibiting E1 decarboxylase activity with 2OA as substrate (*V*_*max*_ 14.2 µmol/min/mg protein, *K*_*m, 2OA*_ 0.2 mM).

**Fig. 1.**
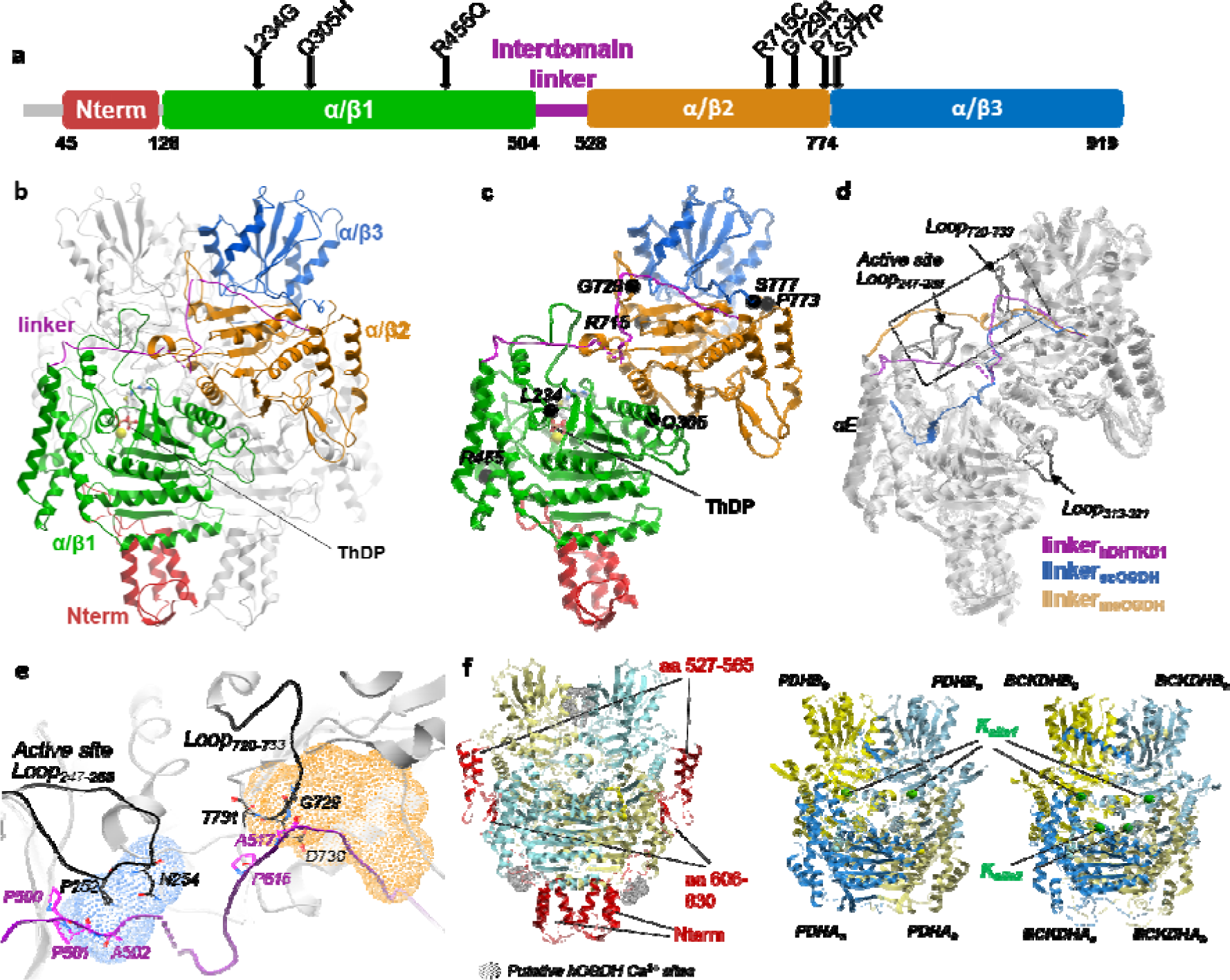
Overall structure of human DHTKD1. (**a**) Schematic of hDHTKD1 domain organisation, indicating disease-causing missense mutations in arrows. (**b**) Structure of the hDHTKD1 homodimer, where one subunit is coloured according to domain organisation from panel a, and the other subunit is coloured grey. ThDP cofactor and Mg^2+^ ion are shown as sticks and spheres respectively. (**c**) Sites of missense mutations are shown as black spheres on one hDHTKD1 protomer (same view as panel b). (**d**) Structural superposition of hDHTKD1, ecOGDH (PDB 2JGD) and msOGDH (PDB 6R2B) highlighting their different inter-domain linkers (coloured purple, blue and light brown respectively). (**e**) Magnified view of the dotted box in panel d, showing how the hDHTKD1 inter-domain linker (purple) packs against two loop regions (black). Binding positions for the allosteric effectors AMP in the ecOGDH structure (PDB 2JGD) and acetyl-CoA in the msOGDH structure (PDB 2Y0P) are shown as blue and brown meshes. (**f**) Overall architecture of the hDHTKD1 homodimer (left), human (PDHA-PDHB)_2_ heterotetramer (middle, PDB 1NI4), and human (BCKDHA-BCKDHB)_2_ heterotetramer (right, PDB 1DTW) are shown. hDHTKD1 homodimer has a larger volume due to the loop insertions coloured red.

The crystal structure of hDHTKD1_45-919_ is determined to 1.9 Å resolution by molecular replacement, using the *E. coli* OGDH structure (PDB 2JGD, 38% sequence identity, [27]) as search template (Table 1). The asymmetric unit contains two DHTKD1 protomers (A and B; Fig. 1b) arranged as an intertwined obligate homodimer in a similar manner as OGDH, burying a large 5600 Å^2^ (18%) area of monomeric accessible surface at the dimer interface. This crystal homodimer is consistent with small-angle x-ray scattering (SAXS) analysis of hDHTKD1_45-919_ protein in solution, with the theoretical scattering curve of the dimer displaying a good fit to experimental data (χ^2^ of 3.8, Suppl. Fig. 2).

**Table 1:** Summary of diffraction and refinement statistics.

|  |  |
| --- | --- |
| **Data Collection** |  |
| Space group | P1 |
| Cell dimensions |  |
| a, b, c (Å) | 78.55 81.22 86.9 |
| $\alpha, \beta, \gamma$ (°) | 63.43 76.96 72.06 |
| Wavelength (Å) | 0.976 |
| Resolution (Å)<sup>a</sup> | 46.01 – 1.87 (1.937 – 1.87) |
| Rmerge (%) | 0.10 (0.44) |
| I/σI | 4.96 (1.04) |
| CC1/2 | 0.987 (0.645) |
| Completeness | 96.45 (91.94) |
| Multiplicity | Multiplicity 1.8 (1.5) |
| **Refinement** |  |
| Resolution (Å) | 1.9 |
| No. reflections | 144961 (13819) |
| Rwork/Rfree | 0.19/0.22 |
| No. atoms |  |
| Protein | 13331 |
| Ligand/ion | 55 |
| Water | 1066 |
| Bfactor (Å<sup>2</sup>) |  |
| Protein | 18.8 |
| Solvent | 28.07 |
| R.m.s. deviations |  |
| Bond lengths (Å) | 0.007 |
| Bond angles (°) | 0.84 |
| Ramachandran plot |  |
| Most favoured (% , residues) | 96.9 |
| Allowed (% , residues) | 2.87 |
| Disallowed (% , residues) | 0.23 |

<sup>a</sup>Numbers in parentheses represent data in the highest resolution shell.

Our DHTKD1 structural model (Fig. 1c) consists of residues 53-915 from both chains, with the exception that no electron density was observed for two surface exposed loop regions (aa 274-275_chain A_/274-277_chain B_ and aa 502-508_chain A_/505-508_chain B_). DHTKD1 is structurally composed of an N-terminal helical bundle (aa 53-127) followed by three α/β domains (α/β1: aa 129-496, Pfam PF00676; α/β2: aa 528-788, PF02779; α/β3: aa 789-915, PF16870). These four structural regions assemble into two halves, inter-connected by an extended linker (aa 497-527) that threads along the protein surface (Fig. 1b,c).

### Structural comparison of DHTKD1 with other E1 enzymes

Expectedly, a Dali search [31] reveals that the closest structural homologue to hDHTKD1 is msOGDH (z-score 55.7, rmsd 1.9 Å, 38% sequence identity) and ecOGDH (z 50.3, 1.8 Å, 40%). The main structural divergence is found in their interdomain linkers, which traverse the respective protein surface via different trajectories (Fig. 1d). Importantly, the hDHTKD1 linker packs against two loop regions that are longer than the equivalents in ecOGDH and msOGDH (Fig. 1e). These include the ‘active site loop’ (aa 247-258), and a DHTKD1-unique region (aa 720-733) identified as ‘Δ3’ in [11] (Suppl. Fig. 3). The path traversed by the hDHTKD1 linker also overlaps with the binding sites for the allosteric activators of ecOGDH (acetyl-CoA, [27]) and msOGDH (AMP, [28]) revealed from their structures (Fig. 1e, meshes). These allosteric sites are likely not present in DHTKD1 structure, due to low sequence conservation in the neighbourhood.

To a lesser degree, hDHTKD1 is also structurally homologous to the E1 enzymes of human PDH and BCKDH (also known as 2-oxoisovalerate dehydrogenase) (z 30, 3.0-3.5 Å, 16%, [25,26]), which are heterotetramers built from two copies of two subunits (Fig. 1f). This contrasts with DHTKD1, OGDH and presumably OGDHL which are homodimers. PDH and BCKDH form a more compact shape, lacking several surface insertions to the α/β core that are unique to the DHTKD1/OGDH/OGDHL subgroup. These include the helical bundle at the N-terminus, and the β- and helical hairpins (aa 527-565, 606-630) within the α/β2 domain (Fig. 1f, red ribbons). PDH and BCKDH structures also contain K^+^ binding sites that play a role in enzymatic regulation (Fig. 1f, green spheres). We did not observe any difference density that suggests metal binding in the equivalent region of DHTKD1. Metal-dependent regulation is also featured in mammalian OGDH enzymes [32,33], mediated by Ca^2+^ binding motifs unique to the OGDH N-terminus and a region equivalent to the DHTKD1 Δ3 [34]. Again, these motifs are not present in prokaryotic OGDHs, or DHTKD1.

### The DHTKD1 active site favours 2OA as substrate

Each DHTKD1 subunit in the crystal homodimer is bound with a ThDP cofactor (Fig. 1b,c), at a site formed from both subunits (Suppl. Fig. 4). The ThDP pyrophosphate moiety binds to the α/β1 domain of one subunit, the pyrimidine ring binds to the α/β2 of the other subunit, while the central thiazolium ring sits between subunits. The ThDP binding residues are highly conserved among OGDH and E1 homologues (Suppl. Fig. 3). These include Asp333 and Asn366 which bridge the ThDP pyrophosphates with the Mg^2+^ ion. Also, Leu290 acts as a hydrophobic wedge to form the characteristic V-shaped conformation of ThDP, bringing the N4’ amino group of pyrimidine ring into close proximity (3.08 Å) with the C2 proton of thiazolium ring. Essential for catalysis, Glu640 triggers a proton relay to activate the cofactor into a reactive ylide (Suppl. Fig. 4). Reaction then ensues via a nucleophilic attack by the ThDP ylide on the substrate keto carbon of the substrate, forming a pre-decarboxylation intermediate that is in turn decarboxylated into an enamine-like ThDP adduct.

Compared to an *apo* E1 structure such as that of ecOGDH, our ThDP-bound DHTKD1 structure highlights 4 loop segments in the active site that undergo disorder-to-order transition during cofactor binding (Fig. 2a). Using nomenclature from ref [11] (Suppl. Fig. 4), these include: ‘Region 1’ (aa 187-195), which contributes Tyr190 to the substrate binding site); the active site loop (aa 247-258), with different length and sequence from OGDHs; ‘loop 1’ (aa 366-383), which contributes the L_368_GY_370_ motif to bind the ThDP pyrophosphate; and ‘loop 2’ (aa 434-445), which contributes residues to engage with E2 enzyme for acyltransfer (e.g. His435). The conformations seen in our *holo* structure are similar to those of msOGDH structures bound with the post-decarboxylation cofactor conjugates [29] (Fig. 2b).

**Fig. 2.**
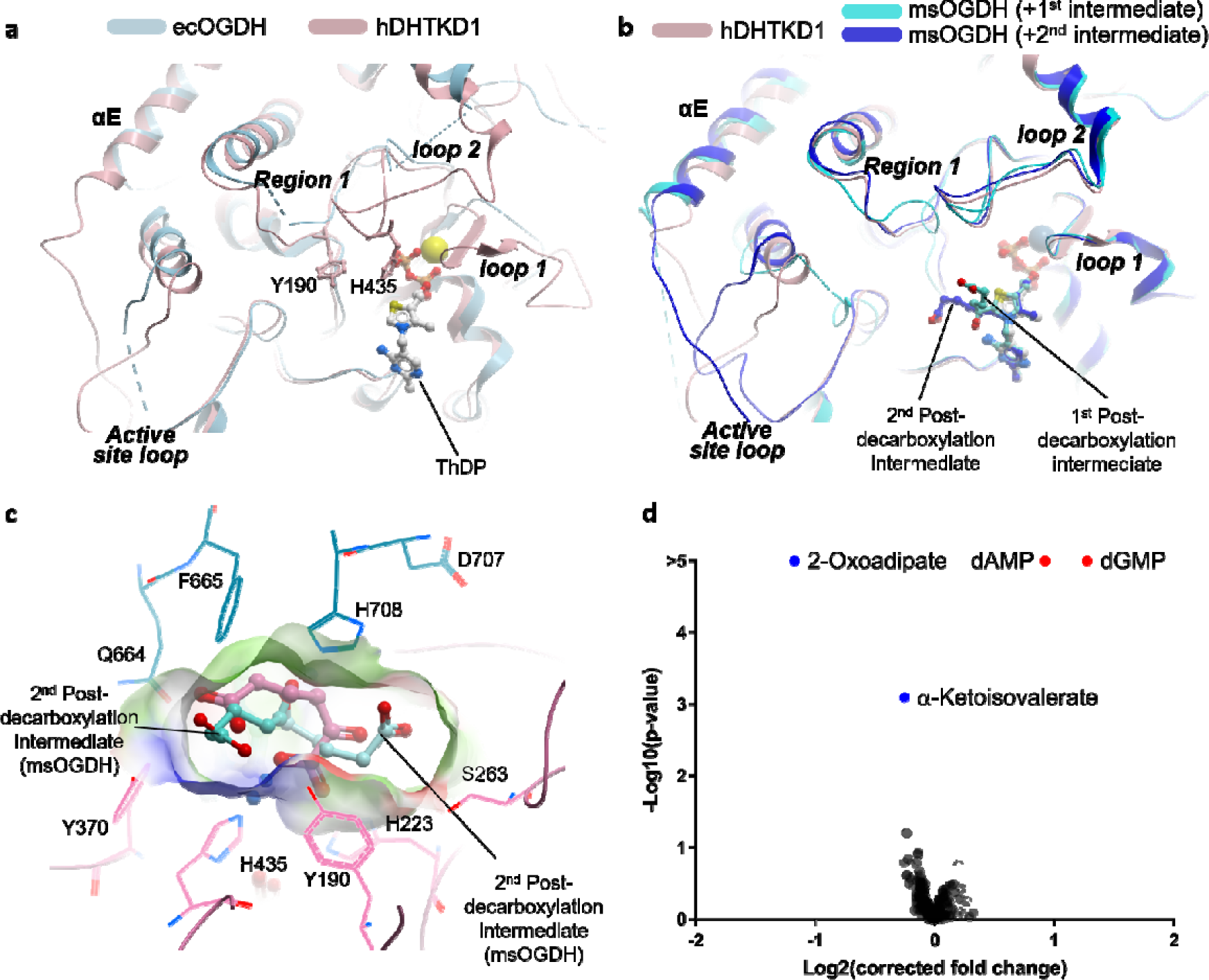
Substrate binding pocket of human DHTKD1. (**a**) View of the active site from the hDHTKD1 structure (pink ribbon) bound with ThDP (sticks, white carbon atoms) and Mg^2+^ (yellow), overlaid with the ecOGDH structure in the apo form (blue ribbon; PDB 2JGD), to highlight structural differences in various loop regions. (**b**) Same hDHTKD1 active site view as in panel a, overlaid with msOGDH structures in complex with the first post-decarboxylation intermediate (cyan ribbon for protein, sticks with cyan carbon atoms for ligand; PDB 3ZHT), and second post-decarboxylation (blue ribbon for protein, sticks with blue carbon atoms for ligand; PDB 3ZHU). (**c**) hDHTKD1 substrate pocket is lined by residues from both subunits of the homodimer (pink and blue lines). Overlaid in this pocket is the putative 2OA ligand (pink sticks) modelled in our density (Suppl. Fig. 5), and the post-decarboxylation intermediates (cyan sticks) bound to the msOGDH structure. (**d**) The hDHTKD1-metabolite interactome as determined by MIDAS. hDHTKD1 significantly depleted (•) 2OA and α-oxoisovalerate and enriched (•) dAMP and dGMP. The cut-off for significance was p < 0.05, q < 0.1.

In one x-ray dataset, we observed omit map electron density at one active site of the homodimer that is not accounted for by the cofactor or any component of the crystallisation condition (Suppl. Fig. 5a). This density is adjacent to but disjoint from the ThDP cofactor, at a location partly overlapping the two conformations of post-decarboxylation intermediate seen in the msOGDH structures (PDB: 3ZHT, 3ZHU; [29]) (Fig. 2b). The size of this density feature can accommodate a C6 ligand such as 2-oxoadipate (2OA) without covalent linkage to ThDP. Although the observed ligand, likely co-purified with the protein, did not undergo enzymatic turnover, the keto carbon can be placed at 3.5 Å from the ThDP thiazolium C2 and hence compatible with the nucleophilic attack and subsequent decarboxylation (Suppl. Fig. 5b). While in good agreement with omit map, the 2OA model was not included in the deposited structure, in light of no further experimental evidence of its presence.

DHTKD1 and OGDH overlap to some extent in their *in vitro* reactivity towards 2OG and 2OA [13,35]. For example, soaking msOGDH crystals with 2OA and 2OG both yielded similar post-decarboxylation intermediates [29]. Nevertheless, hDHTKD1 turns over 2OA with 40-fold catalytic efficiency over 2OG [13]. The hDHTKD1 active site reveals several amino acids poised to interact with the substrate, but vary in sequence with OGDH orthologues. Two of them involve substitution to more polar residues i.e. Tyr190 (from *Phe*_OGDH_) and Tyr370 (*Phe*_OGDH_), while the other two to less bulky residues i.e Ser263 (*Tyr*_OGDH_) and Asp707 (*Glu*_OGDH_) (Suppl. Fig. 3). Overlaying the two msOGDH post-decarboxylation intermediates (first and second conformers, sticks in Fig. 2b; [29]) onto the hDHTKD1 substrate pocket clearly explained how these substituted amino acids can stabilise catalytic intermediates generated from the longer 2OA substrate (Fig. 2c). The 2OA terminal carboxyl group from the first conformer in msOGDH (PDB 3ZHT) can be sandwiched between hDHTKD1 Tyr190 (*Phe*_OGDH_) and Tyr370 (*Phe*_OGDH_) to form polar interactions, while hDHTKD1 Ser263 (Tyr_OGDH_) and Asp707 (*Glu*_OGDH_) increase pocket volume to accommodate the terminal carboxyl group from the second conformer in msOGDH (PDB 3ZHU). It remains to be determined whether the post-decarboxylation intermediate of hDHTKD1 also exists in dual conformation. Suffice it to rationalise that the DHTKD1 substrate pocket is engineered to accommodate the slightly larger and more polar 2OA substrate, providing a structural basis for its superior catalytic efficiency over 2OG.

### DHTKD1 preferentially interacts with 2OA in solution

We further explored the substrate preference of hDHTKD1 by mapping its metabolite interactome using MIDAS, a mass spectrometry-based equilibrium dialysis approach [36]. From a screening library of 412 human metabolites, 2OA was observed as the most significant (p < 4.33×10^−54^, q < 2.59×10^−51^) interaction with hDHTKD1_45-919_ in the presence of ThDP and Mg^2+^ (Fig. 2d). Furthermore, 2OA had the most negative Log_2_[corrected fold change] value (−1.17), suggesting that hDHTKD1 enzymatically processed 2OA during the MIDAS screening. Relative to 2OA, the 2OG interaction with hDHTKD1 was not significant (p < 0.17, q < 0.74) and had a relatively small negative Log_2_[corrected fold change] value (−0.28). The higher confidence and fold change observed for 2OA, relative to 2OG, are in complete agreement with the substrate preference of hDHTKD1.

α-Oxoisovalerate, the primary product of valine degradation by branched-chain-amino-acid aminotransferases, was the second most significant metabolite (p < 8.00×10^−4^, q < 2.06×10^−2^) and had second most negative Log_2_[corrected fold change] value (−0.25), suggesting hDHTKD1 could interact with and may also enzymatically process α-oxoisovalerate. The deoxypurine monophosphates, dAMP and dGMP, had significant (p < 4.47×10^−11^, q < 4.84E-09 and p < 5.13×10^−18^, q < 1.07×10^−15^) positive Log2(corrected fold change) values (0.92 and 1.27), suggesting binding to hDHTKD1. These results support observations that purine nucleotides functionally regulate eukaryotic OGDHc [33,37], and perhaps OADHc. Further experiments are warranted to understand the functional relevance of α-oxoisovalerate and nucleotide monophosphates on DHTKD1 activity.

### DHTKD1 and DLST form direct interactions

There is literature evidence that the E1 and E2 components of 2-oxoacid dehydrogenase complexes interact directly as a binary subcomplex, in the absence of E3. For some E1 enzymes such as OGDH and PDH, the N-terminus is known to be important for the direct interaction with E2 [38,39] and E3 [40], although this region is notably different for DHTKD1. For example, the hDHTKD1 precursor encodes a mere 50-aa segment before the first α-helix of the structure, while the hOGDH equivalent region is longer (121 aa) and contains two DLST-binding motifs [38] not preserved in DHTKD1 (Suppl. Fig. 1b). This suggests that the manner in which DHTKD1 and OGDH (E1) interact with DLST (E2) could be different.

Human DLST as a precursor protein is structurally composed of (Fig. 3a): the mitochondrial target sequence (aa 1-67), the N-terminal single lipoyl domain (aa 68-154) to which a dihydrolipoyl moiety is covalently attached through a lysine residue (Lys110), the C-terminal catalytic domain responsible for the multimeric assembly and harbouring the acyl□transferase active site (aa 211-453), and the flexible inter-domain linker (aa 155-210). We opted to reconstitute the DHTKD1-DLST binary complex by co-expressing His-tagged hDHTKD1_45-919_ and hDLST_68-453_ in *E. coli* followed by affinity chromatography. Untagged hDLST_68-453_ was found to co-purify with His-tagged hDHTKD1_45-919_ immobilised on Ni affinity resin (Fig. 3b), and to similar extent with hDHTKD1_1-919_ (Fig. 3c) and hDHTKD1_23-919_ (Fig. 3d, Suppl. Fig. 6). Hence the hDHTKD1 N-terminal 45 aa, not present in our structural model and replacing the DLST-binding motifs mapped for hOGDH, does not play a role in the DHTKD1-DLST interaction. Size exclusion chromatography (SEC) using an analytical Superose 6 Increase column eluted the complex at V_e_ = 10.4 ml, as compared to hDHTKD1_45-919_ protein alone which eluted later at V_e_ = 16.3 ml (Fig. 3d). Our attempts to mix the binary DHTKD1-DLST complex with purified DLD did not yield a stable three-way complex in SEC, as was the case shown for OGDHc previously [38].

**Fig. 3.**
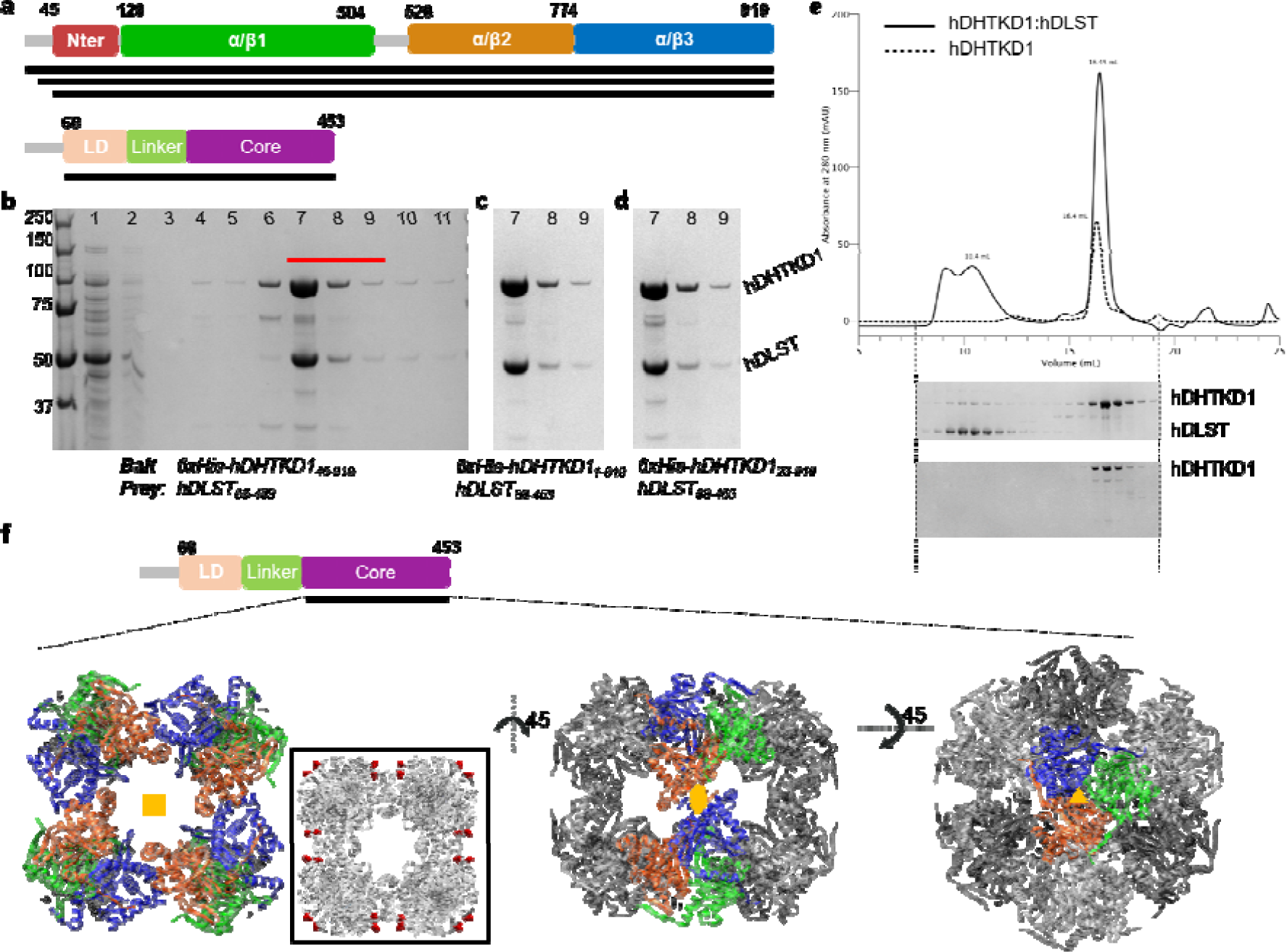
Interaction studies of human DHTKD1 and DLST. (**a**) Constructs of hDHTKD1 and hDLST used in the pulldown experiments. (**b-d**) SDS-PAGE showing affinity pulldown of untagged hDLST_68-453_ from His_6_-tagged hDHTKD1_45-919_ (b), hDHTKD1_1-919_ (c), and hDHTKD1_23-919_ (d). The original uncropped SDS-PAGE gels are shown in Suppl. Fig. 6. For panel b, lanes loaded are: 1, flow-through; 2-6, wash fractions of increasing imidazole concentration; 7-11, elution fractions with 250 mM imidazole. SDS-PAGE gel fragments from panels c and d show only the first three elution fractions i.e. lanes 7, 8, 9 (equivalent to lanes marked red in panel b). (**e**) Chromatogram and SDS-PAGE from size exclusion chromatography runs of hDHTKD1_45-919_ protein alone (dashed line), and of hDHTKD1_45-919_ co-expressed with hDLST_68-453_. Elution volumes for the complex peak and hDHTKD1-alone peak are shown. (**f**) EM Map of the hDLST 24-mer catalytic core, overlaid with a humanized model of *E. coli* DLST (PDB 1SCZ). The three views show how the trimer building block (monomers shown as blue, orange and green ribbons) are assembled into the 24-mer core via four-fold (left), two-fold (middle) and three-fold (right) symmetry axes respectively. Inset of the left view shows how the first residue (aa 219, red spheres) from each of the 24 hDLST catalytic cores are distributed at the surface of the cube structure.

Similar DHTKD1-DLST complex can also be formed by co-expression in the baculo Sf9 cells. When expressed alone in Sf9, the hDLST_68-453_ protein is highly prone to degradation, with a significant proportion fragmenting into two halves (Suppl. Fig. 7a). When the DHTKD1 and DLST proteins are co-expressed, hDHTKD1_45-919_ co-purified in SEC together with both the hDLST_68-453_ intact protein and the C-terminal fragment (containing the catalytic core), while the N-terminal fragment (containing the lipoyl domain and linker) was not part of this complex (Suppl. Fig. 7b). This suggests that the DLST N-terminal fragment alone is not sufficient to interact with DHTKD1, although this DLST region was previously mapped to be interacting with the binding motifs at the hOGDH-unique N-terminus [38]. Altogether, our data reinforce the notion that DHTKD1 and OGDH interact with DLST differently.

### Insight into complex assembly from cryo-EM and SAXS studies

To provide a structural context for the DHTKD1-DLST interactions, we attempted single particle cryo-electron microscopy (cryo-EM) on the reconstituted binary complexes co-expressed in *E. coli* and Sf9 cells. Electron micrographs showed the characteristic cubic cage structures (of approximate dimensions 130 Å x 130 Å x 130 Å), as observed in *E. coli* DLST [41] and other E2 enzymes such as *A. vinelandii* PDH E2 [42] and bovine BCKDH E2 [43]. Upon close inspection, we observed in the *E. coli* co-expressed complex more heterogenous particles, some of which reveal extra density emanating from the cubic core to approximately 10-20 Å (Suppl. Fig. 8a). This was not the case in the Sf9 co-expressed complex, where particles are more homogenous and contain only cubic cages (Suppl. Fig. 8b). To date, we managed to collect a dataset of 33072 particles from the latter sample, and generated a 3D reconstruction at 5 Å resolution (Suppl. Fig. 9). This shows 24 DLST C-terminal catalytic domains assembled as eight trimer building blocks into a cubic cage with 432 symmetry (Fig. 3f). Our EM map allows tracing of a humanised DLST model (aa 219-453 of human DLST) based on the *E. coli* structure (PDB 1E2O; 60% identity) [41].

Considering the sequence conservation, the catalytic cores of *E. coli* and human DLST display essentially identical topology and symmetry along 2-, 3- and 4-fold axes (Fig. 3f). In this assembly, all 24 C-terminal catalytic domains have their first residue (aa 219) exposed to the surface of the core (Fig. 3f, inset), presumably projecting the adjacent inter-domain linker outwards from the core in order to deliver the N-terminal lipoyl domain for engagement with E1 and E3. We reasoned that the additional density protruding from the core in some of our cryo-EM grids (Suppl. Fig. 8a) represent an ordered segment of the linker region. Of the cryo-EM grids imaged thus far, there is unfortunately no discernible density for further regions of DLST (e.g. N-terminal lipoyl domain) or for the DHTKD1 protein. It is likely that the DLST lipoyl domain and much of the linker region are highly dynamic, in agreement with previous attempts to structurally characterize other full-length E2 enzymes such as human PDH E2 [44]. Additionally, the DHTKD1-DLST interaction could be short-lived, as shown for other E1-E2 complexes.

The positioning of 24 lipoyl domains at the exterior of the DLST core implies that they are all potentially available for engagement with E1 and E3. To explore the underlying stoichiometry for the DHTKD1-DLST interaction, we characterised the Sf9 co-expressed binary complex using SEC-SAXS (Suppl. Fig. 10). MW of hDHTKD1_45-919_:hDLST_68-453_ derived from SAXS porod volume is 2.45 MDa, which is in close agreement with a MW of 2.7 MDa calculated for 24x DLST and 16x DHTKD1 protomers, assuming a stoichiometry of one DHTKD1 dimer per DLST trimer building block, as suggested previously for hOGDHc [38]. It remains unknown whether the two active sites within a DHTKD1 dimer are engaged by one or two lipoyl domains. With either possibility, it is apparent that not all 24 lipoyl domains from one DLST core were engaged with DHTKD1 at the same time.

### Structural mapping of disease-causing DHTKD1 mutations

To date 7 *DHTKD1* missense mutations have been reported as the molecular cause for 2-aminoadipic and 2-oxoadipic aciduria. At the protein level, three (p.L234G, p.Q305H, p.R455Q) are located within the α/β1 domain, while the other four (p.R715C, p.G729R, p.P773L, p.S777P) are clustered in the α/β2 domain (Fig. 1a,c). From >100 DHTKD1 and OGDH orthologues surveyed, the aa 715 position is invariantly Arg, while aa positions 305 and 777 are also highly conserved (82% and 93% respectively) (Fig. 4a). None of these residues directly affect the conserved catalytic machinery common to the 2-oxoacid dehydrogenase family.

**Fig. 4.**
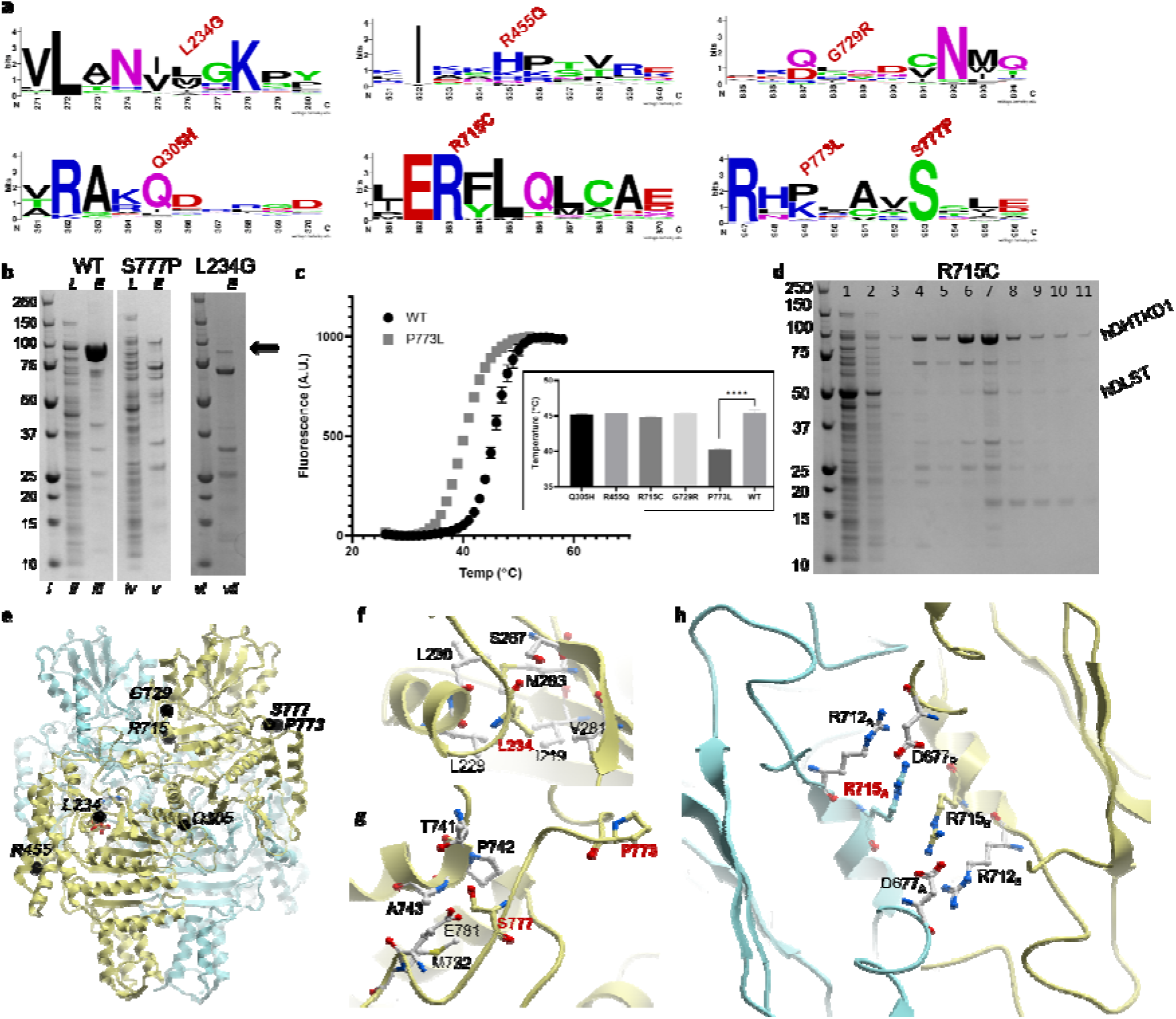
Characterisation of human DHTKD1 disease variants. (**a**) Weblogo diagrams generated from an alignment of >100 DHTKD1 orthologues and homologues, showing sequence conservation for the regions surrounding the seven missense mutation sites. (**b**) Small scale expression and purification for hDHTKD1 wt and variants. SDS-PAGE gel slices showing lysate (*L*) and eluant (*E*) samples from affinity purification of hDHTKD1_45-919_ wt, p.S777P and p.L234G. Position of the hDHTKD1 bands is marked by arrow. Original uncropped gels are shown in Suppl. Fig. 11 (lanes i-v were cropped from one gel, lanes vi-vii were cropped from another). (**c**) DSF melting curves for hDHTKD1_45-919_ wt and p.P773L, with inset showing the derived melting temperature (T_*M*_) values for wt and 5 variants. (**d**) Affinity pulldown of hDLST_68-453_ by immobilised His-tagged hDHTKD1_45-919_ p.R715C variant. SDS-PAGE shown is one of two biological replicates (Suppl. Fig. 13). Lanes are loaded with following samples: 1, flow-through; 2-6, wash fractions of increasing imidazole concentration; 7-11, elution fractions with 250 mM imidazole. (**e**) hDHTKD1 homodimer showing sites of the seven missense mutations (black spheres) on one subunit (yellow ribbon). (**f-h**) Atomic environments surrounding the mutation sites p.L234G (f), p.P773L and p.S777P (g), and p.R715C (h)

To understand their putative biochemical defects, we reconstructed the 7 *DHTKD1* missense mutations recombinantly. All hDHTKD1_45-919_ variants were expressed as soluble protein to similar level as wild-type (wt), with the exception of p.L234G and p.S777P which showed significantly lower yields and propensity to degradation, suggesting these variant proteins are misfolded, compared to wt (Fig. 4b, Suppl. Fig. 11). Leu234 is located at the protein centre ∼20 Å from cofactor site (Fig. 4e), and the p.L234G change introduces a smaller side-chain thereby leaving a cavity at the hydrophobic core (Fig. 4f). Ser777 is partially exposed to the protein surface (Fig. 4e), and the p.S777P change introduces a proline side-chain that likely disrupts hydrogen bonds with neighbouring residues (Fig. 4g).

The remainder 5 variant proteins were isolated, purified, and subjected to thermostability analysis by differential scanning fluorimetry. Four of them (p.Q305H, p.R455Q, p.R715C, p.G729R) demonstrated similar single unfolding-folding transition as hDHTKD1 wt (Fig. 4c, inset). However, p.P773L exhibited a significantly reduced melting temperature (ΔTm = −5.2 °C), suggesting that while expressed as soluble protein this variant is thermally more labile than wt (Fig. 4c). Pro773 forms a bend for the surface exposed loop which connects the α/β2 and α/β3 domains and also harbours the abovementioned Ser777. Replacing Pro773 with Leu likely alters the structural integrity of this loop (Fig. 4h) and could affect protein folding. The observation of two destabilising mutations within this one loop region strongly implicates its importance in the functioning of DHTKD1.

Arg715 is located at the two-fold axis of the homodimer, and together with Arg712 forms a salt bridge network with Asp677 of the opposite subunit (Fig. 4h). Arg712, Arg715 and Asp677 are invariant amino acid positions across DHTKD1 and OGDH orthologues, indicating the importance of this salt bridge network. Arg715 is positioned immediately after ‘loop 3’, a signature motif conserved across all E1 enzymes, including the invariant His708 that is involved in the reductive acyl transfer to E2 (Suppl. Fig. 3) [45]. SEC of the p.R715C variant revealed similar chromatogram profile as wt hDHTKD1_45-919_, indicating intact dimer formation (Suppl. Fig. 12). Nevertheless, when assayed in our co-expression and affinity pull-down, the hDHTKD1_45-919_ p.R715C variant has significantly reduced ability to bind hDLST_68-453_ directly (Fig. 4d, Suppl. Fig. 13), compared to wt (Fig. 3b). Therefore the effect of p.R715C substitution could be transmitted from the dimer interface to engagement with E2 through loop 3. As control, hDHTKD1_45-919_ bearing the p.R455Q or p.P773L substitution (both located at protein exterior) interacts with hDLST_68-453_ to similar extent as wt.

Our data did not reveal any discernible defects on protein stability or interaction with DLST *in vitro* for the p.Q305H, p.R455Q and p.G729R variants, the latter two being found in the majority of reported cases of 2-aminoadipic and 2-oxoadipic aciduria. These results imply that additional functions or unknown binding partners could be involved in the OADHc. Future efforts can be focussed on studying their *in vivo* impact using patient-relevant cells.

## CONCLUSION

DHTKD1 is emerging as a key player in mitochondrial metabolism through its influence in lysine metabolism, energy production, and ROS balance. The structure of DHTKD1 presented here provides the first template for the rational design of DHTKD1 small molecule modulators, to probe the enzyme’s role in these mitochondrial functions and the associated disease states. DHTKD1 exhibits key structural differences from others E1 enzymes particularly OGDH, providing a molecular basis for the subtle difference in substrate specificity and protein-protein interaction despite their close homology. These features would likely be exploited by the DHTKD1-specific modulators.

We have reconstituted the DHTKD1-DLST complex *in vitro*, and demonstrated for the first time that complex formation is disrupted in some disease-causing variants, likely via indirect (e.g. destabilising DHTKD1 to reduce its steady state level) or direct (e.g. altering the binding interface of DHTKD1) mechanisms. These data underscore the importance of DHTKD1 functioning within the context of the OADHc complex. Our Cryo-EM and complementary studies provide insights into how DLST forms a multimeric core to recruit multiple DHTKD1 protomers into this mega assembly. Considering that both DHTKD1 and OGDH recruit DLST as their E2 component, future studies are warranted to explore the existence of a ‘hybrid’ complex in which lipoyl domains from one DLST multimeric core could engage with both DHTKD1 and OGDH at the same time. This could allow crosstalk and regulation between the OADHc and OGDHc complexes for their concerted mitochondrial functions.

## MATERIALS AND METHODS

### Expression and purification of human DHTKD1 and DLST

Site-directed mutations were constructed using the QuikChange mutagenesis kit (Stratagene) and confirmed by sequencing. All primers are available upon request. Wild-type and variant DHTKD1 proteins, as well as all DLST proteins, were expressed in *Escherichia coli* BL21(DE3)R3-Rosetta cells from 1-6 L of Terrific Broth culture. Cell pellets were lysed by sonication and centrifuged at 35,000x *g*. The clarified cell extract was incubated with Ni-NTA resin pre-equilibrated with lysis buffer (50 mM HEPES pH 7.5, 500 mM NaCl, 20 mM Imidazole, 5% Glycerol, 0.5 mM TCEP). The column was washed with 80 ml Binding Buffer (50 mM HEPES pH 7.5, 500 mM NaCl, 5% glycerol, 20 mM Imidazole, 0.5 mM TCEP), 80 ml Wash Buffer (50 mM HEPES pH 7.5, 500 mM NaCl, 5% glycerol, 40 mM Imidazole, 0.5 mM TCEP) and eluted with 15 ml of Elution Buffer (50 mM HEPES pH 7.5, 500 mM NaCl, 5% glycerol, 250 mM Imidazole, 0.5 mM TCEP). The eluant fractions were concentrated to 5 ml and applied to a Superdex 200 16/60 column pre-equilibrated in GF Buffer (50 mM HEPES pH 7.5, 500 mM NaCl, 0.5 mM TCEP, 5% glycerol). Eluted protein fractions were concentrated to 10-15 mg/ml. Lipoylation of DLST proteins was verified by intact mass spectrometry.

### Co-expression of DHTKD1 and DLST

The DHTKD1-DLST complex used in this study was co-expressed in both *E. coli* and insect *Sf9* cells. For *E. coli* co-expression, hDHTKD1_45-919_ was subcloned into the pCDF-LIC vector (incorporating His-tag), and the resultant plasmid was co-transformed with the plasmid encoding untagged hDLST_68-453_ in the pNIC-CT10HStII vector. Co-transformed cultures were grown and protein purification was performed as described above for DHTKD1 alone. For co-expression in insect cells, Sf9 culture was co-infected with one baculovirus expressing His-tagged hDHTKD1_45-919_ and one baculovirus expressing His-tagged hDLST_68-453_.

### Crystallization and structure determination of DHTKD1

Crystals were grown by vapour diffusion method. To crystalize hDHTKD1_45-919_, sitting drops containing 75 nL protein (10 mg/mL) and 75 nL well solution containing 20% (w/v) PEG 3350, 0.1 M bis-tris-propane pH 8.5, 0.2 M sodium formate and 10% (v/v) ethylene glycol were equilibrated at 4°C. Diffraction data were collected at the Diamond Light Source beamline i04, and processed using the CCP4 program suite [46]. hDHTKD1_45-919_ crystallized in the primitive space group P1 with two molecules in the asymmetric unit. The structure was solved by molecular replacement using the program *PHASER* [47] and the *E. coli* OGDH structure (PDB code 2JGD) as search model. The structure was refined using *PHENIX* [48], followed by iterative cycles of model building in *COOT* [49]. Statistics for data collection and refinement are summarized in Table 1.

### DHTKD1 enzyme assay

The enzymatic activity assay was performed in a buffer containing 35 mM potassium phosphate (KH_2_PO_4_), 0.5 mM EDTA, 0.5 mM MgSO_4_, 2 mM 2OA or 2OG, 1 mM ThDP, 5 mM sodium azide (NaN_3_) and 60 µM 2,6-dichlorphenol indophenol (DCPIP), pH 7.4. The activity was determined as a reduction of DCPIP at λ= 610-750 nm, 30°C [50], with and without 2OA or 2OG. The dye DCPIP changes colour from blue to colourless, when being reduced [51]. To obtain *K*_*m*_ and *V*_*max*_ different concentrations of 2OA (0.1 mM, 0.05 mM, 0.1 mM, 0.25 mM, 0.5 mM, 0.75 mM, 1 mM, 2 mM) and no substrate were measured in a 96-well micro titre plate (total well volume 300 µl). The ensuing OD values were plotted on a graph (slope= 1/*V*_*max*_; Y-intercept= *K*_*m*_/*V*_*max*_) to calculate *K*_*m*_ and *V*_*max*_ using the Hanes Woolf plot:

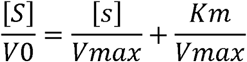

*V*_*0*_= initial velocity; [S]= substrate concentration; *V*_*max*_= maximum velocity.

### Small angle X-ray scattering

SAXS experiments were performed at 0.99 Å wavelength Diamond Light Source at beamline B21 coupled to the appropriate size exclusion column (Harwell, UK) and equipped with Pilatus 2M two-dimensional detector at 4.014 m distance from the sample, 0.005 < q < 0.4 Å^-1^ (q = 4π sin θ/λ, 2θ is the scattering angle). hDHTKD1_45-919_ at 20 mg/ml in 10 mM Hepes-NaOH pH 7.5, 200 mM NaCl, 0.5 mM TCEP and 2% glycerol was applied onto the Shodex KW404-4F column. hDHTKD1_45-919_ co-expressed with hDLST_68-453_ in baculo Sf9 cells at 5 mg/mL in 25 mM Hepes pH 7.5, 200 mM NaCl, 0.5 mM TCEP, 2% Glycerol, 1% sucrose was applied onto the Shodex KW404-4F column. hDLST at 10 mg/ml in 25 mM Hepes pH 7.5, 200 mM NaCl, 0.5 mM TCEP, 2% glycerol, 1% sucrose was applied onto the Shodex 405-4F column.

SAXS measurements were performed at 20 □C, using an exposure time of 3 seconds per frame. SAXS data were processed and analyzed using the ATSAS program package [52] and Scatter (http://www.bioisis.net/scatter). The radius of gyration Rg and forward scattering I(0) were calculated by Guinier approximation. The maximum particle dimension Dmax and P(r) function were evaluated using the program GNOM [53].

### Solution Analysis

Analytical gel filtration was performed on a Superdex 200 Increase 10/300 GL column or Superose 6 Increase 10/300 GL (GE Healthcare) pre-equilibrated with 20 mM HEPES pH 7.5, 150 mM NaCl and 0.5 mM TCEP.

### Differential scanning fluorimetry (DSF)

DSF was performed in a 96-well plate using an Mx3005p RT-PCR machine (Stratagene) with excitation and emission filters of 492 and 610 nm, respectively. Each well (20 µl) consisted of protein (2 mg/ml in 100 mM Hepes pH 7.5, 150 mM NaCl, 5% glycerol), SYPRO-Orange (Invitrogen, diluted 1000-fold of the manufacturer’s stock). Fluorescence intensities were measured from 25 to 96 °C with a ramp rate of 1 °C/min. *T*_m_ was determined by plotting the intensity as a function of temperature and fitting the curve to a Boltzmann equation. Temperature shifts, Δ *T*_m_, were determined as described [54] and final graphs were generated using GraphPad Prism (v.7; Graph-Pad Software). Assays were carried out in technical triplicate.

### MIDAS protein-metabolite screening

Protein-metabolite interaction screening using an updated MIDAS platform was performed similar to Orsak, *et al* [36]. FIA-MS spectra collected from MIDAS protein-metabolite screening was qualitatively and quantitatively processed in SCIEX OS 1.5 software to determine relative metabolite abundance by integrating the mean area under the curve (AUC) across technical triplicates.

### Grid preparation and data acquisition

3□µl of 0.4□mg/ml purified complex sample was applied to the glow-discharged Quantifoil Au R1.2/1.3 grid (Structure Probe), and vitrified using a Vitrobot Mark IV (FEI Company). Cryo grids were loaded into a Glacios transmission electron microscope (ThermoFisher Scientific) operating at 200□keV with a Falcon3 camera. Images were recorded linear mode with pixel size of 0.96□Å and a defocus range of −1 to −3.1□μm with steps of 0.3 μm. Data were collected with a dose rate of 32.52 e/A^2 and images were recorded with a 1s exposure (19 total frames). All details corresponding to individual datasets are summarized in Supplementary Table 2.

### EM data collection and processing

A total of 619 dose-fractioned movies were beam-induced motion correction using MotionCor2 [55] with the dose-weighting option. CTF parameters were determined by ctffind4.1 [56]. The DLST particles were auto-picked using Relion 3.0 [57]. A total of 165739 particles were then extracted with a box size of 344 pixels rescaled to 172 pixels (2x binning). Particles were filtered for homogeneity using rounds of 2D, 3D classification with Relion 3.0 [57]. 33,072 particles where re-extracted un-binned and submitted to 3D auto-refinement with O (octahedral) symmetry imposed resulting in a 5□Å map after post processing (Suppl. Fig. 10a). 3D refinements were started from a 50□Å low-pass filtered version of a 3D class map from previous step. Global resolution was estimated by applying a soft mask around the protein complex density and based on the gold-standard (two halves of data refined independently) FSC□=□0.143 criterion (Suppl. Fig. 10b). Local resolution was calculated with relion 3.0 [57] (Suppl. Fig. 10c).

### EM Model building and refinement

To fit a template to the final map the *E.coli* DLST orthologue structure (PDB 1SCZ) was used. The sequence was humanized and residues truncated to the alpha carbon using Chainsaw [58]. The oligomeric structure was docked into the O symmetry full EM density map using Molrep followed by one round of refinement using refmac5 through CCP-EM [59]. Fig. 5 and Suppl. Fig. 10 were made using Chimera [60].

## ACKNOWLEDGEMENTS

The Structural Genomics Consortium is a registered charity (Number 1097737) that receives funds from AbbVie, Bayer Pharma AG, Boehringer Ingelheim, Canada Foundation for Innovation, Eshelman Institute for Innovation, Genome Canada, Innovative Medicines Initiative (EU/EFPIA) [ULTRA-DD grant no. 115766], Janssen, Merck & Co., Novartis Pharma AG, Ontario Ministry of Economic Development and Innovation, Pfizer, São Paulo Research Foundation-FAPESP, Takeda, and Wellcome Trust [092809/Z/10/Z].

## DATA AVAILABILITY

X-ray coordinates and structure factors were deposited to the Protein Data Bank (accession code 6SY1). Cryo-EM data were deposited to the EMDB (EMD-10556).

## REFERENCES

1. Yeaman SJ (1989) The 2-oxo acid dehydrogenase complexes: recent advances. Biochem J 257 (3):625–632. doi: 10.1042/bj2570625

2. Reed LJ (2001) A trail of research from lipoic acid to alpha-keto acid dehydrogenase complexes. J Biol Chem 276 (42):38329–38336. doi: 10.1074/jbc.R100026200

3. Perham RN (1991) Domains, motifs, and linkers in 2-oxo acid dehydrogenase multienzyme complexes: a paradigm in the design of a multifunctional protein. Biochemistry 30 (35):8501–8512. doi: 10.1021/bi00099a001

4. Jordan F (2003) Current mechanistic understanding of thiamin diphosphate-dependent enzymatic reactions. Nat Prod Rep 20 (2):184–201. doi: 10.1039/b111348h

5. Marrott NL, Marshall JJ, Svergun DI, Crennell SJ, Hough DW, van den Elsen JM, Danson MJ (2014) Why are the 2-oxoacid dehydrogenase complexes so large? Generation of an active trimeric complex. Biochem J 463 (3):405–412. doi: 10.1042/BJ20140359

6. Izard T, Aevarsson A, Allen MD, Westphal AH, Perham RN, de Kok A, Hol WG (1999) Principles of quasi-equivalence and Euclidean geometry govern the assembly of cubic and dodecahedral cores of pyruvate dehydrogenase complexes. Proc Natl Acad Sci U S A 96 (4):1240–1245. doi: 10.1073/pnas.96.4.1240

7. Cohen RD, Pielak GJ (2017) A cell is more than the sum of its (dilute) parts: A brief history of quinary structure. Protein Sci 26 (3):403–413. doi: 10.1002/pro.3092

8. Zhou ZH, McCarthy DB, O’Connor CM, Reed LJ, Stoops JK (2001) The remarkable structural and functional organization of the eukaryotic pyruvate dehydrogenase complexes. Proc Natl Acad Sci U S A 98 (26):14802–14807. doi: 10.1073/pnas.011597698

9. Reed LJ, Hackert ML (1990) Structure-function relationships in dihydrolipoamide acyltransferases. J Biol Chem 265 (16):8971–8974

10. Perham RN, Jones DD, Chauhan HJ, Howard MJ (2002) Substrate channelling in 2-oxo acid dehydrogenase multienzyme complexes. Biochem Soc Trans 30 (2):47–51. doi: 10.1042/

11. Bunik VI, Degtyarev D (2008) Structure-function relationships in the 2-oxo acid dehydrogenase family: substrate-specific signatures and functional predictions for the 2-oxoglutarate dehydrogenase-like proteins. Proteins 71 (2):874–890. doi: 10.1002/prot.21766

12. Araujo WL, Trofimova L, Mkrtchyan G, Steinhauser D, Krall L, Graf A, Fernie AR, Bunik VI (2013) On the role of the mitochondrial 2-oxoglutarate dehydrogenase complex in amino acid metabolism. Amino Acids 44 (2):683–700. doi: 10.1007/s00726-012-1392-x

13. Nemeria NS, Gerfen G, Yang L, Zhang X, Jordan F (2018) Evidence for functional and regulatory cross-talk between the tricarboxylic acid cycle 2-oxoglutarate dehydrogenase complex and 2-oxoadipate dehydrogenase on the l-lysine, l-hydroxylysine and l-tryptophan degradation pathways from studies in vitro. Biochim Biophys Acta Bioenerg 1859 (9):932–939. doi: 10.1016/j.bbabio.2018.05.001

14. Goncalves RL, Bunik VI, Brand MD (2016) Production of superoxide/hydrogen peroxide by the mitochondrial 2-oxoadipate dehydrogenase complex. Free Radic Biol Med 91:247–255. doi: 10.1016/j.freeradbiomed.2015.12.020

15. Nemeria NS, Gerfen G, Nareddy PR, Yang L, Zhang X, Szostak M, Jordan F (2018) The mitochondrial 2-oxoadipate and 2-oxoglutarate dehydrogenase complexes share their E2 and E3 components for their function and both generate reactive oxygen species. Free Radic Biol Med 115:136–145. doi: 10.1016/j.freeradbiomed.2017.11.018

16. Quinlan CL, Goncalves RL, Hey-Mogensen M, Yadava N, Bunik VI, Brand MD (2014) The 2-oxoacid dehydrogenase complexes in mitochondria can produce superoxide/hydrogen peroxide at much higher rates than complex I. J Biol Chem 289 (12):8312–8325. doi: 10.1074/jbc.M113.545301

17. Bunik VI, Brand MD (2018) Generation of superoxide and hydrogen peroxide by side reactions of mitochondrial 2-oxoacid dehydrogenase complexes in isolation and in cells. Biol Chem 399 (5):407–420. doi: 10.1515/hsz-2017-0284

18. Xu W, Zhu H, Gu M, Luo Q, Ding J, Yao Y, Chen F, Wang Z (2013) DHTKD1 is essential for mitochondrial biogenesis and function maintenance. FEBS Lett 587 (21):3587–3592. doi: 10.1016/j.febslet.2013.08.047

19. Sherrill JD, Kc K, Wang X, Wen T, Chamberlin A, Stucke EM, Collins MH, Abonia JP, Peng Y, Wu Q, Putnam PE, Dexheimer PJ, Aronow BJ, Kottyan LC, Kaufman KM, Harley JB, Huang T, Rothenberg ME (2018) Whole-exome sequencing uncovers oxidoreductases DHTKD1 and OGDHL as linkers between mitochondrial dysfunction and eosinophilic esophagitis. JCI Insight 3 (8). doi: 10.1172/jci.insight.99922

20. Fischer MH, Gerritsen T, Opitz JM (1974) Alpha-aminoadipic aciduria, a non-deleterious inborn metabolic defect. Humangenetik 24 (4):265–270. doi: 10.1007/bf00297590

21. Danhauser K, Sauer SW, Haack TB, Wieland T, Staufner C, Graf E, Zschocke J, Strom TM, Traub T, Okun JG, Meitinger T, Hoffmann GF, Prokisch H, Kolker S (2012) DHTKD1 mutations cause 2-aminoadipic and 2-oxoadipic aciduria. Am J Hum Genet 91 (6):1082–1087. doi: 10.1016/j.ajhg.2012.10.006

22. Duran M, Beemer FA, Wadman SK, Wendel U, Janssen B (1984) A patient with alpha-ketoadipic and alpha-aminoadipic aciduria. J Inherit Metab Dis 7 (2):61. doi: 10.1007/bf01805803

23. Hagen J, te Brinke H, Wanders RJ, Knegt AC, Oussoren E, Hoogeboom AJ, Ruijter GJ, Becker D, Schwab KO, Franke I, Duran M, Waterham HR, Sass JO, Houten SM (2015) Genetic basis of alpha-aminoadipic and alpha-ketoadipic aciduria. J Inherit Metab Dis 38 (5):873–879. doi: 10.1007/s10545-015-9841-9

24. Xu WY, Gu MM, Sun LH, Guo WT, Zhu HB, Ma JF, Yuan WT, Kuang Y, Ji BJ, Wu XL, Chen Y, Zhang HX, Sun FT, Huang W, Huang L, Chen SD, Wang ZG (2012) A nonsense mutation in DHTKD1 causes Charcot-Marie-Tooth disease type 2 in a large Chinese pedigree. Am J Hum Genet 91 (6):1088–1094. doi: 10.1016/j.ajhg.2012.09.018

25. A AE, Chuang JL, Wynn RM, Turley S, Chuang DT, Hol WG (2000) Crystal structure of human branched-chain alpha-ketoacid dehydrogenase and the molecular basis of multienzyme complex deficiency in maple syrup urine disease. Structure 8 (3):277–291. doi: 10.1016/s0969-2126(00)00105-2

26. Ciszak EM, Korotchkina LG, Dominiak PM, Sidhu S, Patel MS (2003) Structural basis for flip-flop action of thiamin pyrophosphate-dependent enzymes revealed by human pyruvate dehydrogenase. J Biol Chem 278 (23):21240–21246. doi: 10.1074/jbc.M300339200

27. Frank RA, Price AJ, Northrop FD, Perham RN, Luisi BF (2007) Crystal structure of the E1 component of the Escherichia coli 2-oxoglutarate dehydrogenase multienzyme complex. J Mol Biol 368 (3):639–651. doi: 10.1016/j.jmb.2007.01.080

28. Wagner T, Bellinzoni M, Wehenkel A, O’Hare HM, Alzari PM (2011) Functional plasticity and allosteric regulation of alpha-ketoglutarate decarboxylase in central mycobacterial metabolism. Chem Biol 18 (8):1011–1020. doi: 10.1016/j.chembiol.2011.06.004

29. Wagner T, Barilone N, Alzari PM, Bellinzoni M (2014) A dual conformation of the post-decarboxylation intermediate is associated with distinct enzyme states in mycobacterial KGD (alpha-ketoglutarate decarboxylase). Biochem J 457 (3):425–434. doi: 10.1042/BJ20131142

30. Wagner T, Boyko A, Alzari PM, Bunik VI, Bellinzoni M (2019) Conformational transitions in the active site of mycobacterial 2-oxoglutarate dehydrogenase upon binding phosphonate analogues of 2-oxoglutarate: From a Michaelis-like complex to ThDP adducts. J Struct Biol 208 (2):182–190. doi: 10.1016/j.jsb.2019.08.012

31. Holm L, Sander C (1995) Dali: a network tool for protein structure comparison. Trends Biochem Sci 20 (11):478–480. doi: 10.1016/s0968-0004(00)89105-7

32. Rutter GA, McCormack JG, Midgley PJ, Denton RM (1989) The role of Ca2+ in the hormonal regulation of the activities of pyruvate dehydrogenase and oxoglutarate dehydrogenase complexes. Ann N Y Acad Sci 573:206–217. doi: 10.1111/j.1749-6632.1989.tb14998.x

33. Lawlis VB, Roche TE (1981) Inhibition of bovine kidney alpha-ketoglutarate dehydrogenase complex by reduced nicotinamide adenine dinucleotide in the presence or absence of calcium ion and effect of adenosine 5’-diphosphate on reduced nicotinamide adenine dinucleotide inhibition. Biochemistry 20 (9):2519–2524. doi: 10.1021/bi00512a024

34. Rigden DJ, Galperin MY (2004) The DxDxDG motif for calcium binding: multiple structural contexts and implications for evolution. J Mol Biol 343 (4):971–984. doi: 10.1016/j.jmb.2004.08.077

35. Leandro J, Dodatko T, Aten J, Hendrickson RC, Sanchez R, Yu C, DeVita RJ, Houten SM (2019) DHTKD1 and OGDH display in vivo substrate overlap and form a hybrid ketoacid dehydrogenase complex. bioRxiv

36. Orsak T, Smith TL, Eckert D, Lindsley JE, Borges CR, Rutter J (2012) Revealing the allosterome: systematic identification of metabolite-protein interactions. Biochemistry 51 (1):225–232. doi: 10.1021/bi201313s

37. Craig DW, Wedding RT (1980) Regulation of the 2-oxoglutarate dehydrogenase lipoate succinyltransferase complex from cauliflower by nucleotide. Steady state kinetic studies. J Biol Chem 255 (12):5763–5768

38. Zhou J, Yang L, Ozohanics O, Zhang X, Wang J, Ambrus A, Arjunan P, Brukh R, Nemeria NS, Furey W, Jordan F (2018) A multipronged approach unravels unprecedented protein-protein interactions in the human 2-oxoglutarate dehydrogenase multienzyme complex. J Biol Chem 293 (50):19213–19227. doi: 10.1074/jbc.RA118.005432

39. Park YH, Wei W, Zhou L, Nemeria N, Jordan F (2004) Amino-terminal residues 1-45 of the Escherichia coli pyruvate dehydrogenase complex E1 subunit interact with the E2 subunit and are required for activity of the complex but not for reductive acetylation of the E2 subunit. Biochemistry 43 (44):14037–14046. doi: 10.1021/bi049027b

40. McCartney RG, Rice JE, Sanderson SJ, Bunik V, Lindsay H, Lindsay JG (1998) Subunit interactions in the mammalian alpha-ketoglutarate dehydrogenase complex. Evidence for direct association of the alpha-ketoglutarate dehydrogenase and dihydrolipoamide dehydrogenase components. J Biol Chem 273 (37):24158–24164. doi: 10.1074/jbc.273.37.24158

41. Knapp JE, Mitchell DT, Yazdi MA, Ernst SR, Reed LJ, Hackert ML (1998) Crystal structure of the truncated cubic core component of the Escherichia coli 2-oxoglutarate dehydrogenase multienzyme complex. J Mol Biol 280 (4):655–668. doi: 10.1006/jmbi.1998.1924

42. Mattevi A, Obmolova G, Schulze E, Kalk KH, Westphal AH, de Kok A, Hol WG (1992) Atomic structure of the cubic core of the pyruvate dehydrogenase multienzyme complex. Science 255 (5051):1544–1550. doi: 10.1126/science.1549782

43. Kato M, Wynn RM, Chuang JL, Brautigam CA, Custorio M, Chuang DT (2006) A synchronized substrate-gating mechanism revealed by cubic-core structure of the bovine branched-chain alpha-ketoacid dehydrogenase complex. EMBO J 25 (24):5983–5994. doi: 10.1038/sj.emboj.7601444

44. Yu X, Hiromasa Y, Tsen H, Stoops JK, Roche TE, Zhou ZH (2008) Structures of the human pyruvate dehydrogenase complex cores: a highly conserved catalytic center with flexible N-terminal domains. Structure 16 (1):104–114. doi: 10.1016/j.str.2007.10.024

45. Wynn RM, Machius M, Chuang JL, Li J, Tomchick DR, Chuang DT (2003) Roles of His291-alpha and His146-beta’ in the reductive acylation reaction catalyzed by human branched-chain alpha-ketoacid dehydrogenase: refined phosphorylation loop structure in the active site. J Biol Chem 278 (44):43402–43410. doi: 10.1074/jbc.M306204200

46. CCP4 (1994) The CCP4 suite: programs for protein crystallography. Acta Crystallogr D Biol Crystallogr 50 (Pt 5):760–763

47. McCoy AJ, Grosse-Kunstleve RW, Storoni LC, Read RJ (2005) Likelihood-enhanced fast translation functions. Acta Crystallogr D Biol Crystallogr 61 (Pt 4):458–464

48. Adams PD, Afonine PV, Bunkoczi G, Chen VB, Davis IW, Echols N, Headd JJ, Hung LW, Kapral GJ, Grosse-Kunstleve RW, McCoy AJ, Moriarty NW, Oeffner R, Read RJ, Richardson DC, Richardson JS, Terwilliger TC, Zwart PH (2010) PHENIX: a comprehensive Python-based system for macromolecular structure solution. Acta Crystallogr D Biol Crystallogr 66 (Pt 2):213–221. doi: 10.1107/S0907444909052925

49. Emsley P, Cowtan K (2004) Coot: model-building tools for molecular graphics. Acta Crystallogr D Biol Crystallogr 60 (Pt 12 Pt 1):2126–2132

50. Sauer SW, Okun JG, Schwab MA, Crnic LR, Hoffmann GF, Goodman SI, Koeller DM, Kolker S (2005) Bioenergetics in glutaryl-coenzyme A dehydrogenase deficiency: a role for glutaryl-coenzyme A. J Biol Chem 280 (23):21830–21836. doi: 10.1074/jbc.M502845200

51. VanderJagt DJ, Garry PJ, Hunt WC (1986) Ascorbate in plasma as measured by liquid chromatography and by dichlorophenolindophenol colorimetry. Clin Chem 32 (6):1004–1006

52. Petoukhov MV, Franke D, Shkumatov AV, Tria G, Kikhney AG, Gajda M, Gorba C, Mertens HD, Konarev PV, Svergun DI (2012) New developments in the ATSAS program package for small-angle scattering data analysis. J Appl Crystallogr 45 (Pt 2):342–350. doi: 10.1107/S0021889812007662

53. Svergun DI (1992) Determination of the regularization parameter in indirect-transform methods using perceptual criteria. J App Cryst 25:495–503. doi: https://doi.org/10.1107/S0021889892001663

54. Niesen FH, Berglund H, Vedadi M (2007) The use of differential scanning fluorimetry to detect ligand interactions that promote protein stability. Nature protocols 2 (9):2212–2221. doi: 10.1038/nprot.2007.321

55. Zheng SQ, Palovcak E, Armache JP, Verba KA, Cheng Y, Agard DA (2017) MotionCor2: anisotropic correction of beam-induced motion for improved cryo-electron microscopy. Nat Methods 14 (4):331–332. doi: 10.1038/nmeth.4193

56. Rohou A, Grigorieff N (2015) CTFFIND4: Fast and accurate defocus estimation from electron micrographs. J Struct Biol 192 (2):216–221. doi: 10.1016/j.jsb.2015.08.008

57. Scheres SH, Chen S (2012) Prevention of overfitting in cryo-EM structure determination. Nat Methods 9 (9):853–854. doi: 10.1038/nmeth.2115

58. Stein N (2008) CHAINSAW: a program for mutating pdb files used as templates in molecular replacement. Journal of Applied Crystallography 41 (3):641–643. doi: 10.1107/S0021889808006985

59. Burnley T, Palmer CM, Winn M (2017) Recent developments in the CCP-EM software suite. Acta Crystallogr D Struct Biol 73 (Pt 6):469–477. doi: 10.1107/S2059798317007859

60. Pettersen EF, Goddard TD, Huang CC, Couch GS, Greenblatt DM, Meng EC, Ferrin TE (2004) UCSF Chimera--a visualization system for exploratory research and analysis. J Comput Chem 25 (13):1605–1612. doi: 10.1002/jcc.20084

