## Supplementary Material for "Crystal structure and interaction studies of human DHTKD1 provide insight into a mitochondrial megacomplex in lysine catabolism"

**Supplementary Information**

Supplementary Table S1

Supplementary Fig. S1-13

**Supplementary Table S1** Parameters for cryo-EM data collection and 3D reconstruction

|  | **DLST (**EMD-10556) |
| --- | --- |
| **Data Collection** |  |
| Microscope | Glacios |
| Voltage (keV) | 200 |
| Nominal Magnification | 150,000 x |
| Electron exposure (e/Å^2) | 32.52 |
| Dose rate (e/pixel/sec) | 31.22 |
| Camera | Falcon3 |
| Fractions (no.) | 19 |
| Exposure (sec) | 1 |
| Pixel size (Å) | 0.96 |
| Defocus range (μm) | -1 to -3.1 |
| Micrographs collected (no.) | 619 |
| Final refined particles (no.) | 33,072 |
| **Reconstruction** |  |
| Symmetry imposed | O |
| Resolution (global) (Å) FSC 0.143 | 5 |
| Applied B-factor (Å^2) | -427.7 |

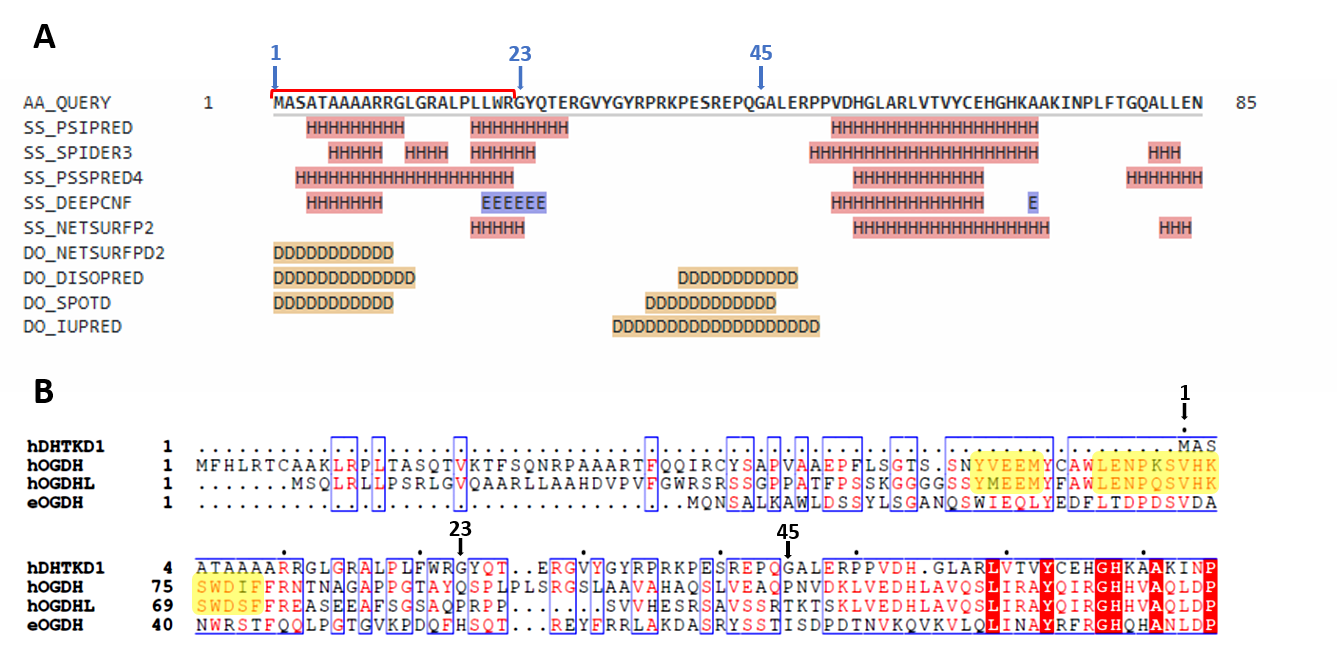

**Supplementary Fig. S1 The N-terminus of human DHTKD1.** (**A**) Secondary structure and intrinsic disorder prediction for the human DHTKD1 N-terminal 88 aa, from 9 different servers (H, helice; E, strand, D, disordered). The putative mitochondrial target sequence is bracketed (red). We have expressed DHTKD1 constructs that begin at residues 1, 23 or 45 (arrows). (**B**) Sequence-based alignment of the N-terminus from human DHTKD1, OGDH, OGDHL and *E. coli* OGDH. Motifs from OGDH (also conserved in OGDHL) that were mapped to interact with DLST (ref 36 from main text) are shaded yellow.

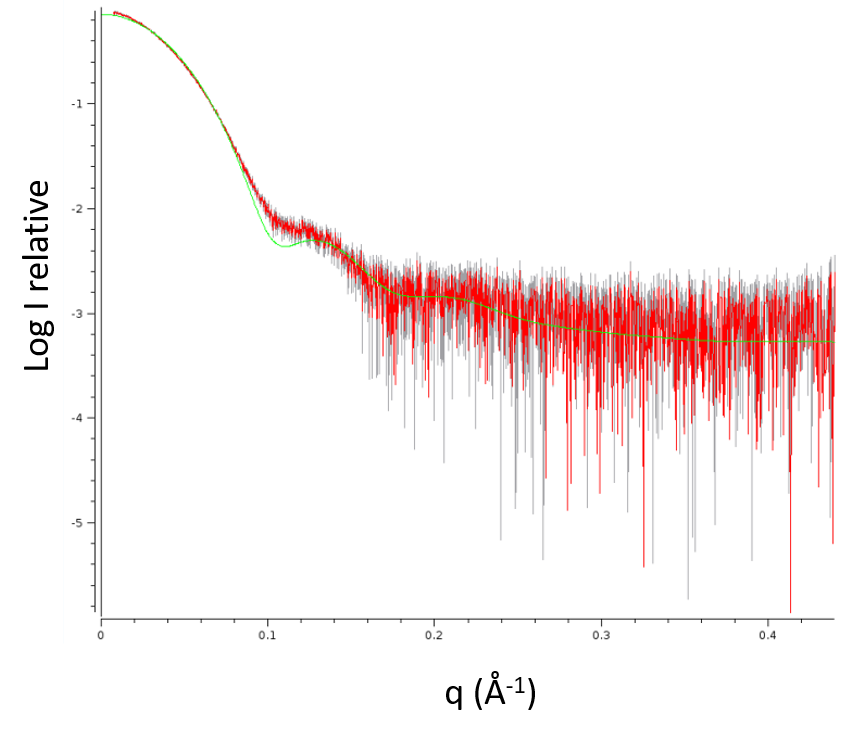

**Supplementary Fig. S2 SAXS analysis of hDHTKD1_45-919_.** Overlay of experimental scattering profile (red) and theoretical scattering curve of DHTKD1 crystal dimer (green). A Chi2 fit of 3.8 was determined by CRYSOL [1].

1. Svergun D, Barberato C, Koch MHJ (1995) CRYSOL - a Program to Evaluate X-ray Solution Scattering of Biological Macromolecules from Atomic Coordinates. Journal of Applied Crystallography 28 (6):768-773. doi:doi:10.1107/S0021889895007047

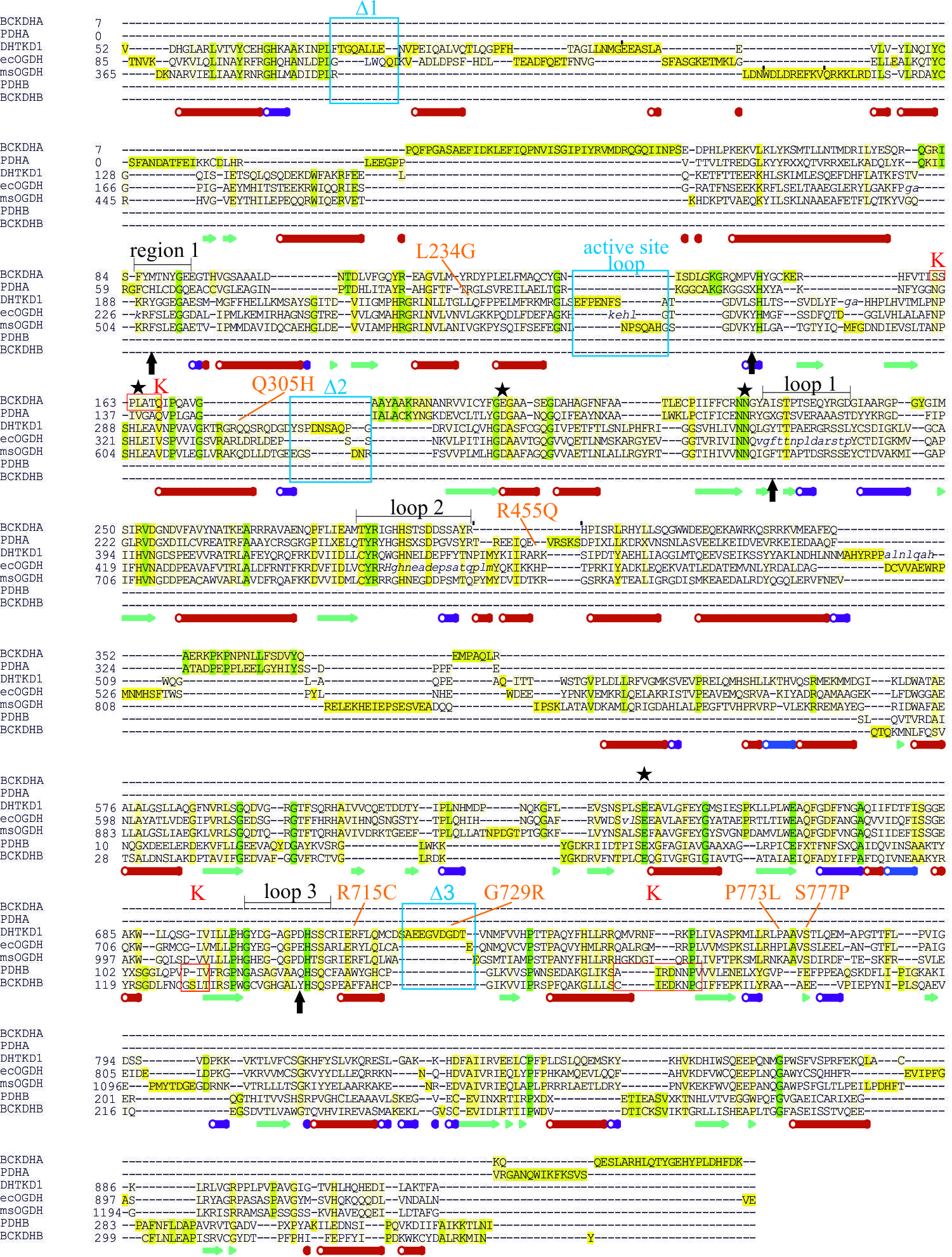

**Supplementary Fig. S3 Structure-based sequence alignment of representative 2-oxoacid dehydrogenases.** Aligned structures using ICM Pro (www.molsoft.com) include human DHTKD1 (this study), human human branched-chain α-ketoacid dehydrogenase subunits BCKDHA, BCKDHB (PDB 1dtw_a, 1dtw_b), human pyruvate dehydrogenase subunits PDHA, PDHB (1ni4_a, 1ni4_b), *E. coli* 2-oxoglutarate dehydrogenase ecOGDH (2jgd), *M. Smegmatis* α-ketoglutarate decarboxylase msOGDH (2y0p). Residues that were not modelled in crystal structures are shown in italicized lowercase. Secondary structures for DHTKD1 are shown at the bottom. Annotated features shown on the alignment include: sites of missense mutations for DHTKD1 (orange fonts), DHTKD1-unique loop regions (blue boxes and font), 4 amino acid positions at the DHTKD1 active site that are distinct from OGDH (black arrows), catalytic residues (black stars) and loop regions (black brackets and font) conserved across the 2-oxoacid dehydrogenase family, and K^+^ ion binding sites from human PDH and BCKDH structures (red boxes and font).

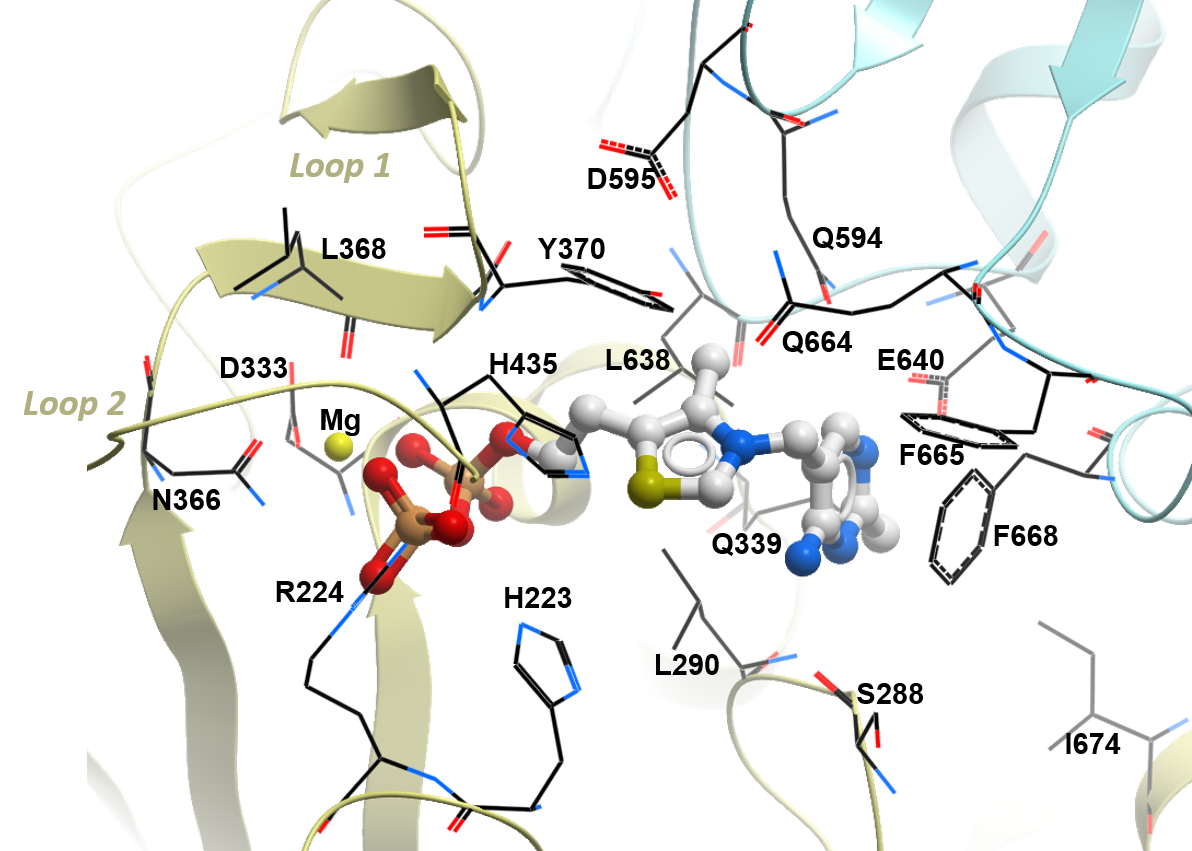

**Supplementary Fig. S4 Binding site of the ThDP cofactor.** ThDP (ball-and-stick) binds to a pocket formed by residues (lines) from both subunits of the DHTKD1 homodimer (yellow and cyan ribbon). Mg^2+^ ion is shown as yellow sphere.

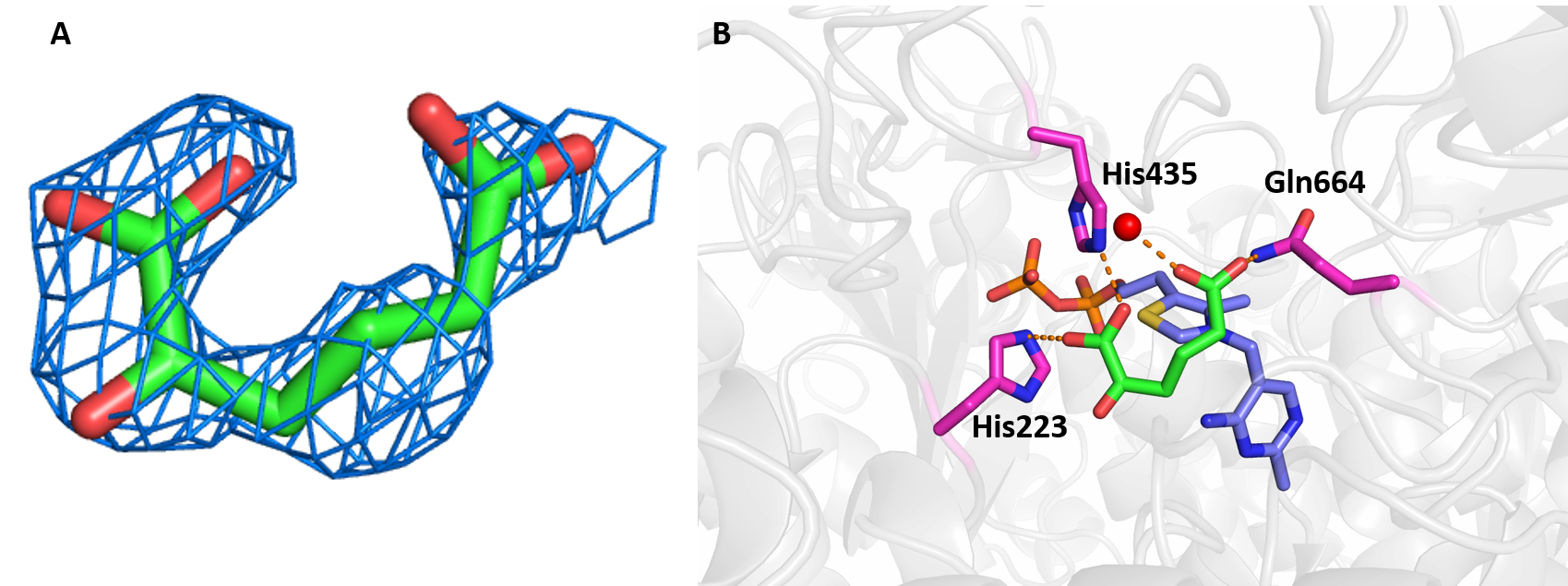

**Supplementary Fig. S5 Putative 2-oxoadipate at the active site.** (**A**) Simulated-annealing 2Fo-Fc composite omit map of hDHTKD1 contoured at 1 σ and displaying the density at the active site, into which a molecule of 2-oxoadipate was modelled. (**B**) Binding environment of the modelled ligand, showing that one carboxyl group of the ligand forms hydrogen bonds with Gln664 and a water molecule, while the other carboxyl group forms hydrogen bonds with His435 and His223. Shown in stick representation are: the modelled ligand with green carbon atoms, ThDP cofactor with lilac carbon atoms, and protein residues with magenta carbon atoms.

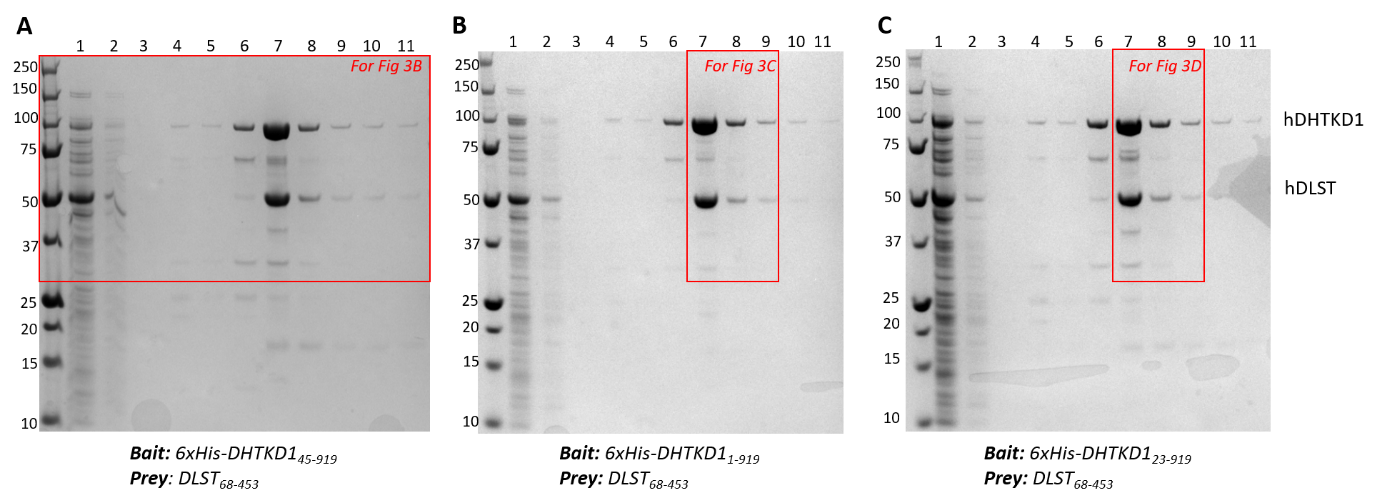

**Supplementary Fig. S6 Affinity pull-down of DLST by immobilised His-tagged DHTKD1.** Recombinant untagged hDLST_68-453_ co-eluted with His_6_-tagged hDHTKD1_45-919_ (**A**), hDHTKD1_1-919_ (**B**), and hDHTKD1_23-919_ (**C**). Full SDS-PAGE gels are shown here. Areas of the gels excised for display in Fig. 3B, C, D of the main text are boxed. Lanes are loaded with following samples from Ni-affinity chromatography: 1, flow-through; 2-6, wash fractions of increasing imidazole concentration; 7-11, elution fractions with 250 mM imidazole.

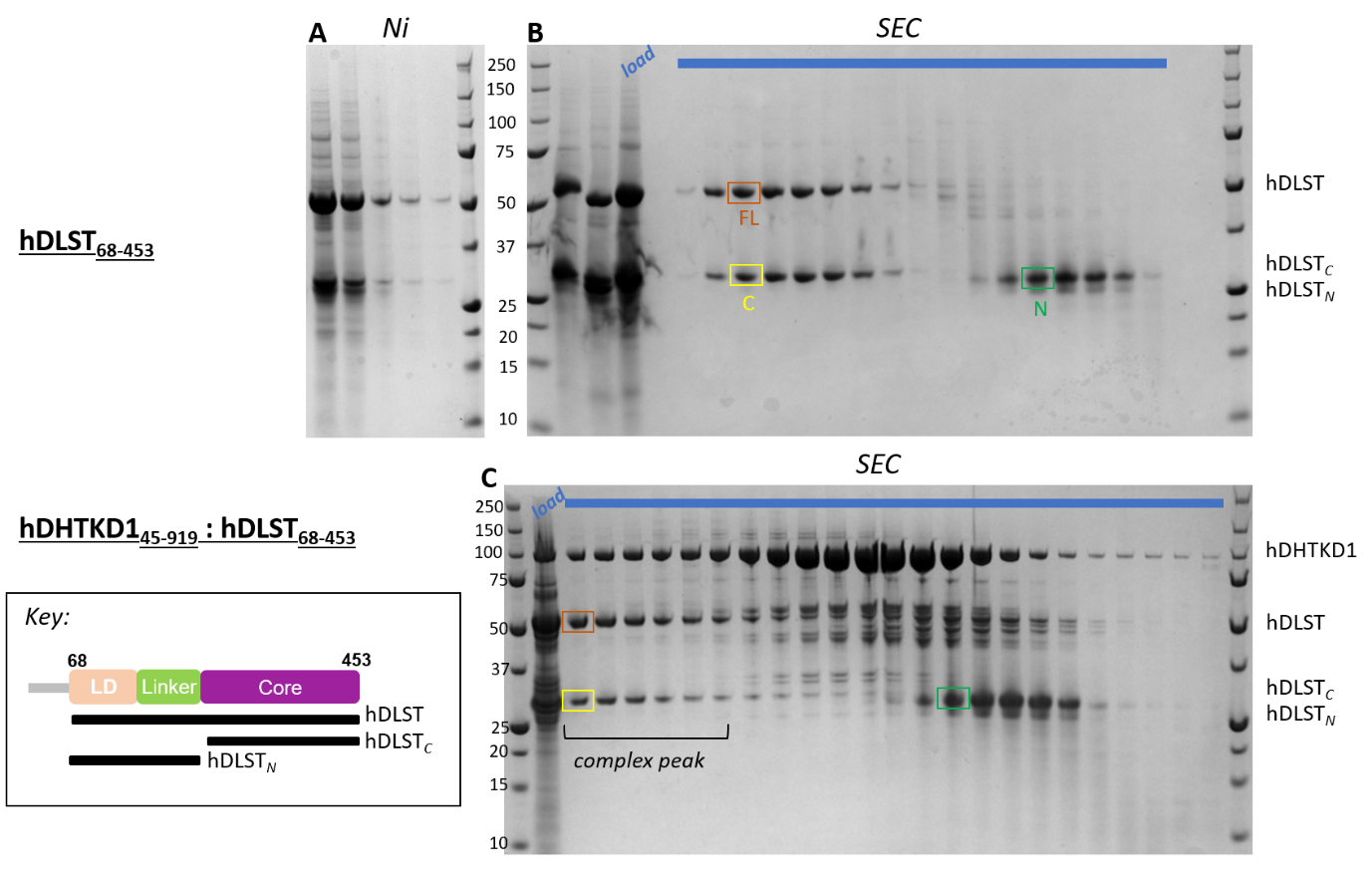

**Supplementary Fig. S7 Purification of human DLST expressed from baculo Sf9 cells.** (**A**) Elution fractions from Ni affinity purification of hDLST_68-453_. (**B**) Elution fractions (blue bar) from size exclusion chromatography (HiLoad 16/600 Superdex 200) of the hDLST_68-453_ sample partially purified from Ni affinity chromatography (as in panel A). Bands in boxes labelled FL, N, C have been verified by tryptic-MS/MS to contain full-length hDLST_48-653_, and the N- and C-terminal halves. (**C**) Elution fractions (blue bar) from size exclusion chromatography (XK 16/70 Superose 6 prep grade) of hDLST_68-453_ co-expressed with hDHTKD1_45-919_ in insect Sf9 cells. Peak fractions from the DHTKD1-containing complex contain full-length as well as the C-terminal half of DLST, but not its N-terminal half.

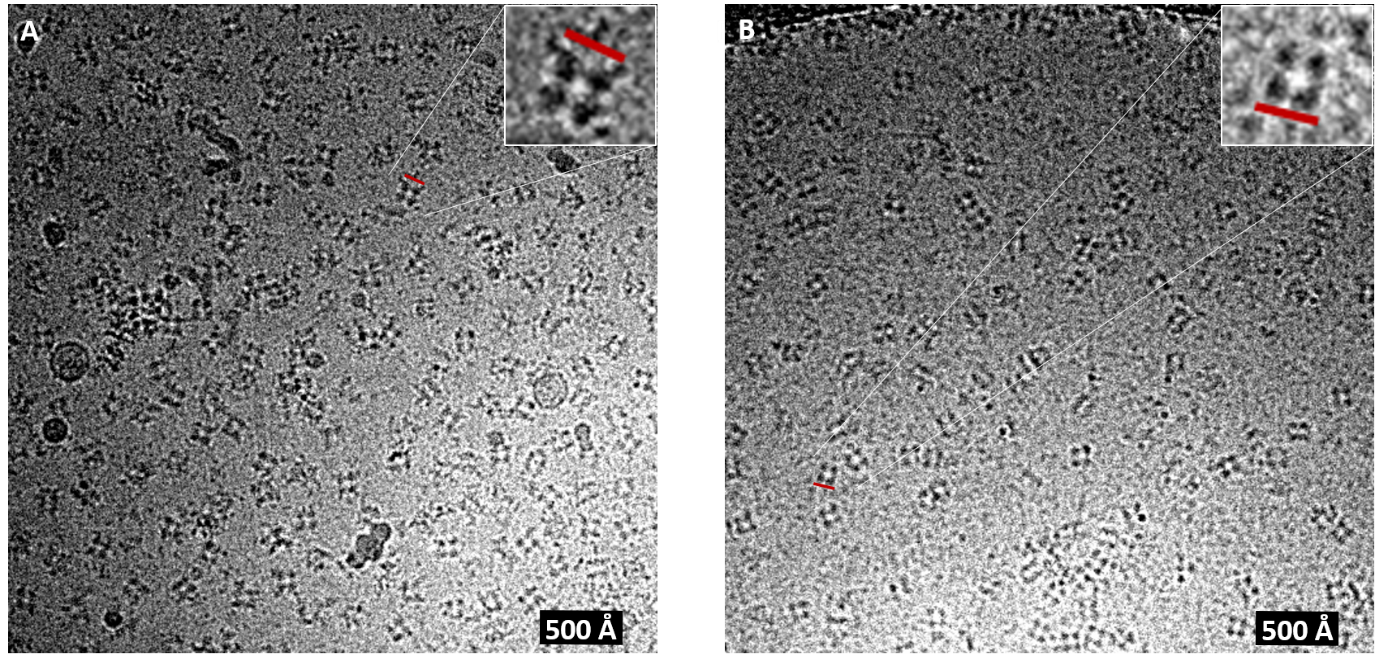

**Supplementary Fig. S8 Electron Micrographs representing two different samples of the DHTKD1-DLST complex.** (**A**) The *E. coli* co-expressed complex reveals heterogenous particles with additional density located at the corners of the DLST cubes. (**B**) The Sf9 co-expressed complex reveals homogenous density of the DLST core catalytic domain only. Red bars indicate a length of approximately 130 Å.

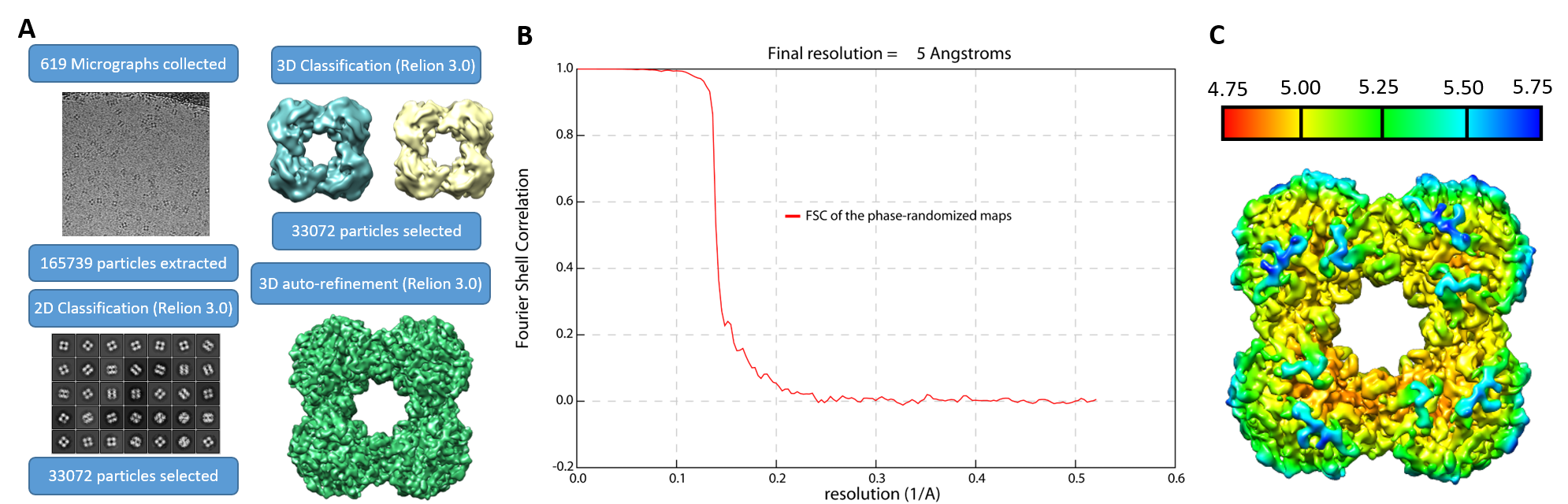

**Supplementary Fig. S9 EM data processing. (A)** Schematic of the processing workflow that resulted in the final 3D reconstruction of the human DLST cube at 5 Å. (**B)** Gold standard FSC curve of phase randomised map showing global resolution of 5 Å. (**C)** Local resolution of the maps estimated using Relion 3.0, scaled between 4.75 and 5.75 Å.

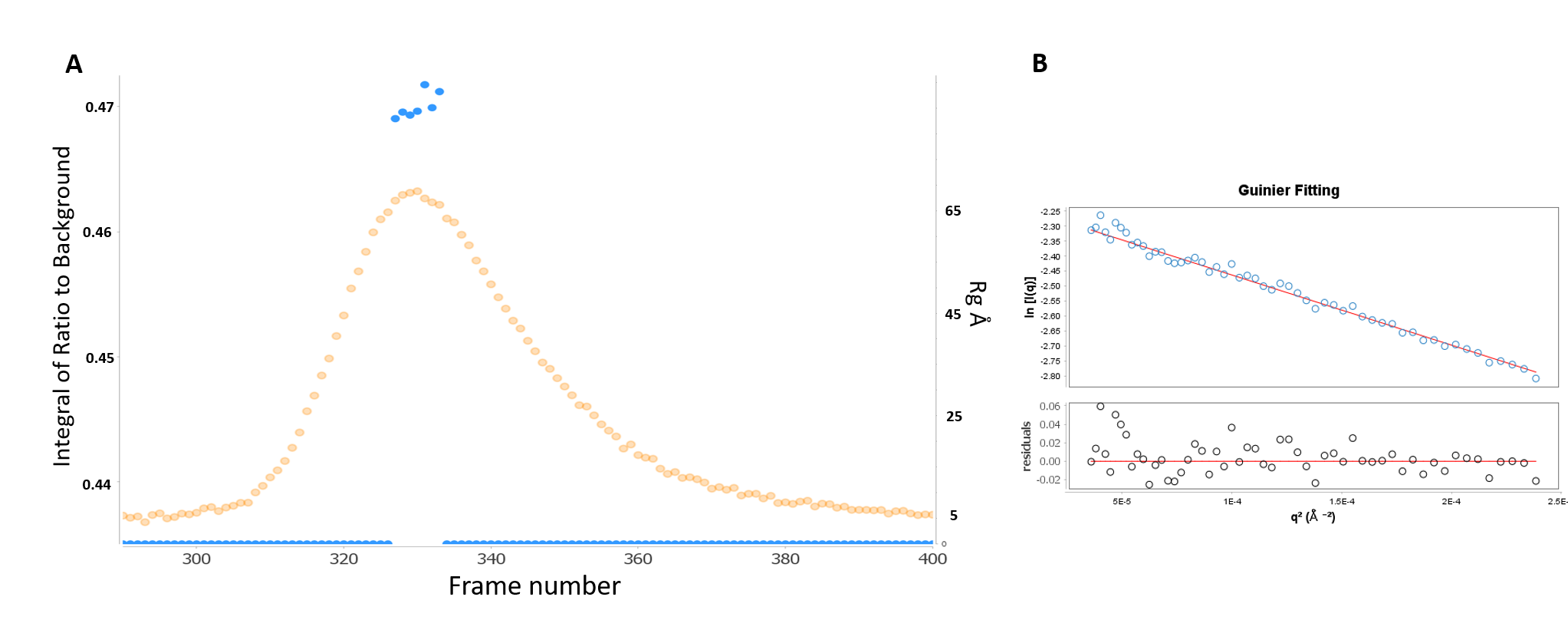

**Supplementary Fig. S10 SEC-SAXS analysis of DHTKD1-DLST complex.** (**A**) Analysis of the radius of gyration (Rg) over elution peak. The Rg (blue spheres) and Integral of Ratio to Background (yellow spheres) are plotted against recorded frames in SEC-SAXS profiles for the hDHTKD1_45-919_:hDLST_68-453_ complex co-expressed in Sf9 cells. The frames used for further analysis are 328-332. Exposure of 3 seconds per frame was recorded. (**B**) Rg value derived from the Guinier plot is 83.2 Å.

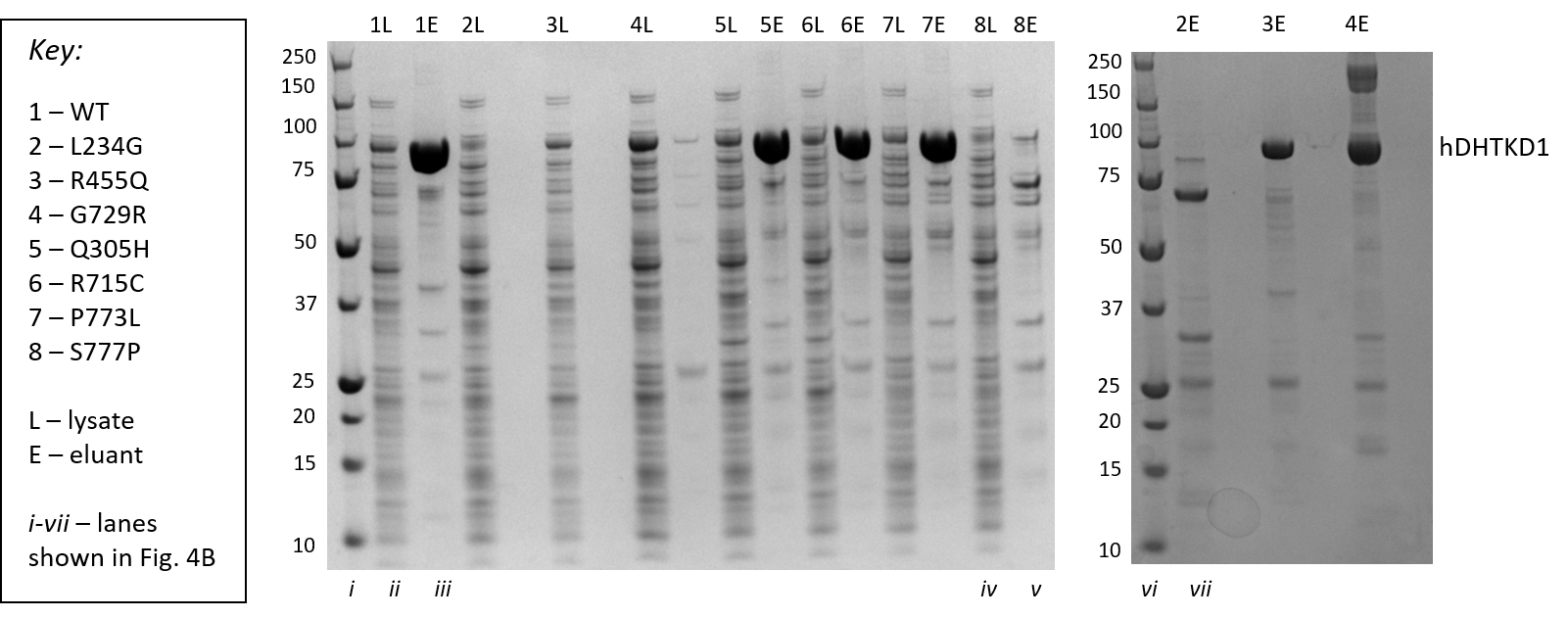

**Supplementary Fig. S11 Small scale expression and purification test for DHTKD1 wt and variants.** SDS-PAGE of lysate (L) and eluant (E) fractions from affinity purification of hDHTKD1_45-919_ wt (1) and missense variants (2-8) are shown. Lanes excised for display in Fig. 4B are labelled in roman numerals i-vii.

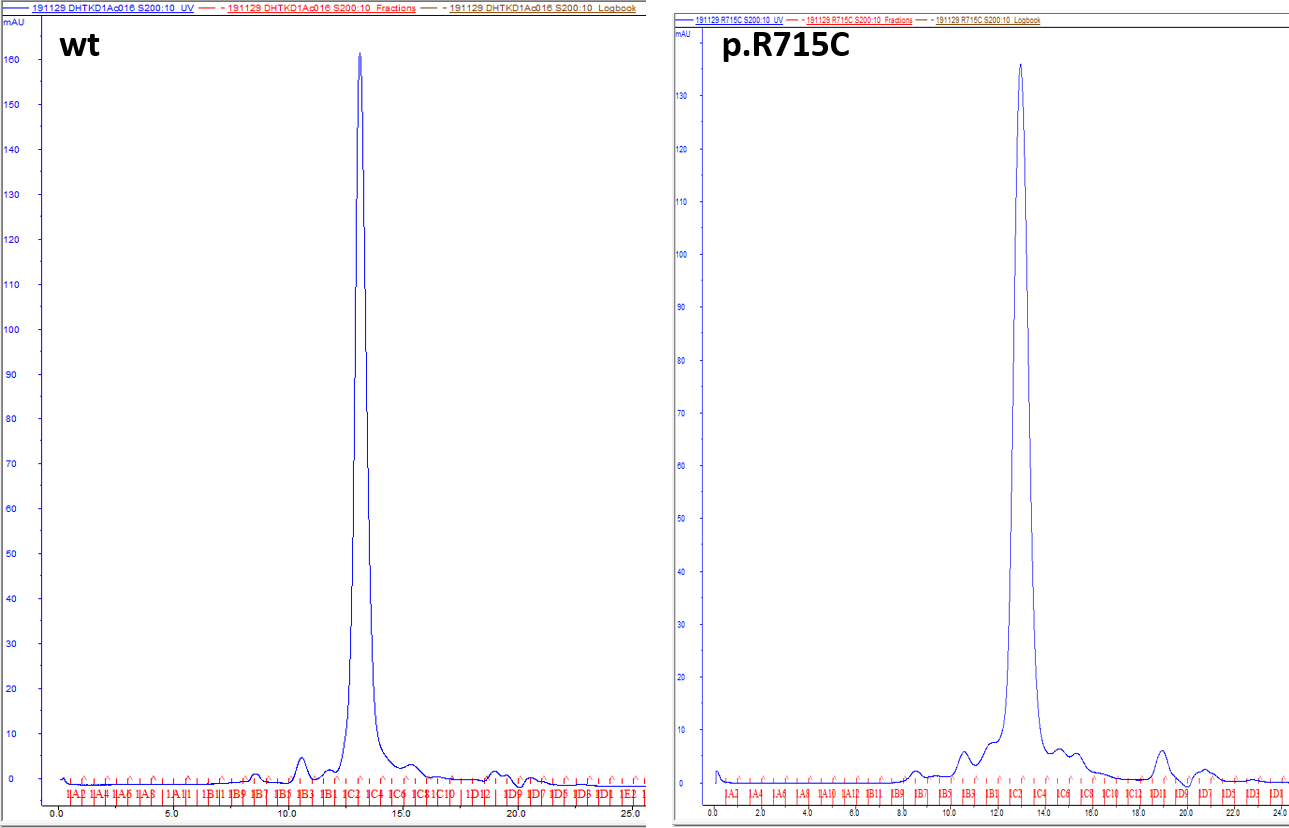

**Supplementary Fig. S12 Size exclusion chromatography of DHTKD1 wt and p.R715C variant.** Chromatograms from Superdex 200 Increase 10/300 GL column are shown, showing that both wt and variant proteins behave similarly in SEC, eluting at the same volume.

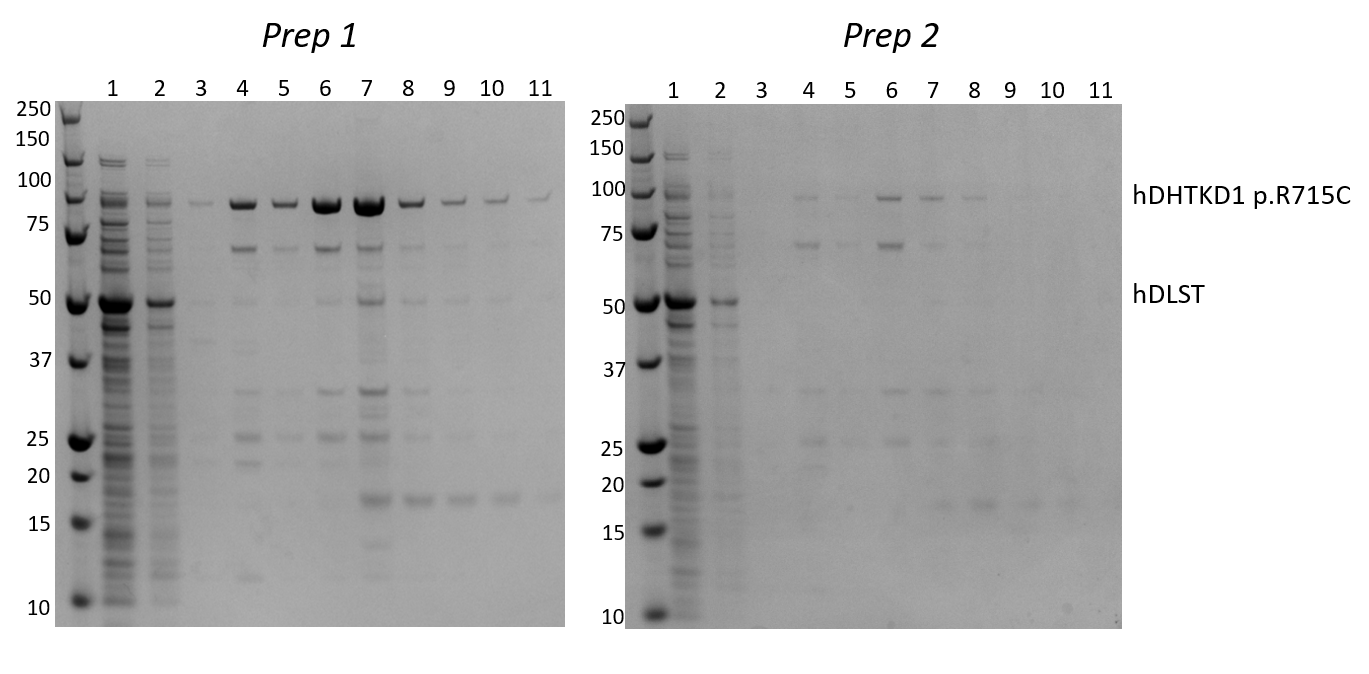

**Supplementary Fig. S13 Affinity pulldown of DLST by immobilised His-tagged DHTKD1 p.R715C variant.** SDS-PAGE are shown from two experimental replicates of different biological preparations. Lanes are loaded with following samples from Ni affinity chromatography: 1, flow-through; 2-6, wash fractions of increasing imidazole concentration; 7-11, elution fractions with 250 mM imidazole. SDS-PAGE for prep 1 is also shown in Fig. 4D.
